# Benchmarking DNA Isolation Methods for Marine Metagenomics Studies

**DOI:** 10.1101/2023.07.25.550485

**Authors:** Alina Demkina, Darya Slonova, Viktor Mamontov, Olga Konovalova, Daria Yurikova, Vladimir Rogozhin, Vera Belova, Dmitriy Korostin, Dmitry Sutormin, Konstantin Severinov, Artem Isaev

## Abstract

Metagenomics is a powerful tool to study marine microbial communities. However, obtaining high-quality environmental DNA suitable for downstream sequencing applications is a challenging task. The quality and quantity of isolated DNA heavily depend on the choice of purification procedure and the type of sample. Selection of an appropriate DNA isolation method for a new type of material often entails a lengthy trial and error process. Further, each DNA purification approach introduces biases and thus affects the composition of the studied community. To account for these problems and biases, we systematically investigated efficiency of DNA purification from three types of samples (water, sea sediment, and digestive tract of a model invertebrate *Magallana gigas*) with eight commercially available microbial DNA isolation kits. For each kit-sample combination we measured the quantity of purified DNA, extent of DNA fragmentation, the presence of PCR-inhibiting contaminants, admixture of eukaryotic DNA, alpha-diversity, and reproducibility of the resulting community composition based on 16S rRNA amplicons sequencing. Additionally, we determined a “kitome”, e.g., a set of contaminating taxa inherent for each type of purification kit used. The resulting matrix of evaluated parameters allows one to select the best DNA purification procedure for a given type of sample.

## Introduction

The Global Ocean is the planet’s largest ecosystem. Yet, biodiversity of Earth’s oceans remains understudied, partly due to the low accessibility of specific niches, such as ocean floor sediments or Arctic regions^1, 2^. The problem is particularly poignant for prokaryotic communities, given that majority of bacteria are unculturable or require very specific growth conditions^3^. In recent years, metagenomics, e.g., sequencing of total DNA isolated directly from collected samples allowed to estimate the real diversity and abundance of environmental microorganisms bypassing the isolation and cultivation steps^4, 5^. Ambitious endeavors, such as the TARA Project^6, 7^, Earth Microbiome Project^8^ or SEA-PHAGES^9^ aim to comprehensively describe microbial or viral diversity across Earth’s marine and terrestrial ecosystems using large-scale metagenomic sequencing. Our project “Atlas of Microbial Communities of the Russian Federation” is conceived to cover one of the largest “white spots” on the Earth’s microbial communities sampling map – the Arctic Ocean.

Several approaches to study the taxonomic and genetic diversity of microbial communities have been developed. 16S rRNA gene hypervariable regions amplicon sequencing allows one to directly assign taxonomy to sequenced DNA fragments and estimate alpha- and beta-diversity, i.e., the diversity of taxa within a sample or between different samples^10^. However, amplicon sequencing introduces biases^11^, does not distinguish close strains^11^ and, more importantly, does not provide information about genes other than 16S rRNA genes. In contrast, by sequencing all DNA fragments found within a sample, short-read shotgun metagenomic sequencing (on Illumina or BGI platforms) allows one, with sufficient quality of input DNA and depth of sequencing, to reconstruct larger genomic fragments (contigs) or even complete bacterial genomes and identify functional gene clusters^12^. In combination with long-read sequencing (using Oxford Nanopore or Pacific Biosciences technologies), one can perform hybrid assembly, which significantly increases the length of contigs but requires higher quality (reduced fragmentation) and quantity of input DNA^13^.

Purification of high-quality metagenomic DNA from natural sources faces a variety of challenges. Sample handling, the choice of DNA purification method, and even homogenization technique significantly affects the resulting community composition^14, 15^. In this work, we processed three types of samples: fresh water, ocean sediments, and digestive system of a model marine invertebrate – Pacific oyster (*Magallana gigas* also known as *Crassostrea gigas*). To select the best strategy for microbial DNA isolation from each type of sample we systematically evaluated available commercial DNA purification kits, and estimated the quality of resulting DNA, it’s applicability for the 16S rRNA amplicon, short-read and long-read sequencing, and compared resulting communities’ composition (Fig. 1). Our goal was to select DNA purification procedures that optimize each of the following parameters:

- DNA quantity, the most evident, yet the most important characteristics, especially for Oxford Nanopore platform that, according to manufacturer recommendations, requires at least 1 μg of DNA (although in some reports this quantity was lowered to just 1 ng^16^. Obtaining sufficient quantities of DNA is also particularly important for Arctic Ocean samples given comparatively low numbers of prokaryotic cells in cold waters^17^.
- DNA purity and the presence of contaminants that can significantly impact the efficiency of PCR and the quality of sequencing libraries^18^. A well-known example of such contaminant is humic acid that often co-purifies with DNA from soil samples and inhibits PCR in concentrations starting from 10 ng/μL^19, 20^. PCR is also sensitive to the presence of chelating agents that sequester Mg^2+^ and to a plethora of organic compounds^18^. Such contaminations can be roughly estimated by UV adsorption at 230 nm, while co-purifying proteins can be detected by UV adsorption at 280 nm.
- DNA fragmentation. Extensive DNA fragmentation can be a consequence of poor sample storage, mechanical shearing during purification, or enzymatic degradation by nucleases released at the stage of cell homogenization^21^. The parameter is especially important for long-read sequencing as it determines the lengths of resulting reads.
- Admixture of eukaryotic DNA. This problem is relevant for shotgun sequencing of DNA from symbiotic microbial communities. Unless specific measures are taken, the proportion of microbial reads can be below a few percent^22^. Although the presence of non-microbial DNA seemingly should not affect 16S rRNA amplicon sequencing, incorrect estimation of the amount of microbial DNA in the sample can lead to PCR amplification biases caused by target DNA underrepresentation.
- Reagent and laboratory contamination with microbial DNA (“kitome” and “splashome”, respectively). This is one of the major problems that precludes correct analysis of microbial communities in sparsely-populated environments, since the signal from contaminating DNA can be higher than that from DNA of interest. Primarily, avoidance of contaminating DNA is achieved by complying with aseptic practices at all stages of sample collection and DNA purification, as well as by sterilization and decontamination of equipment and laboratory space^23^. Although bioinformatic decontamination procedures can be applied at the stage of data analysis, complete removal of non-sample specific sequences is impossible, as the *a priori* community composition is not known^24^. To estimate the signal from contaminating DNA, negative control samples are processed and sequenced in parallel with experimental samples. An unavoidable and often neglected source of DNA contamination is the DNA purification kit itself, as even miniscule quantities of microbial DNA in the kit solutions can be detected during sequencing, and each commercial kit can be described in the terms of the “kitome”, e.g., a set of contaminating taxa it contributes^25, 26^.
- Method-specific biases in the composition of microbial taxa. Another major source of biases in metagenomic analysis is the varied efficiency of DNA purification for different microbial groups. Given the same staring material, different DNA isolation methods will result in vastly different quantities and proportions of group-specific DNA molecules^27, 28^. The major source of this bias is the efficiency of cell lysis. Many gram-positive bacteria or specific groups that form endospores or dormant cells (such as *Firmicutes*, *Actinobacteria*, *Cyanobacteria*, *Myxococcales* and *Azotobacteraceae*) resist homogenization and lysis by convenient methods^29^. At the same time, biases in water samples can be associated with filter retention efficiency that could vary for different cells^30^.

**Fig. 1.**
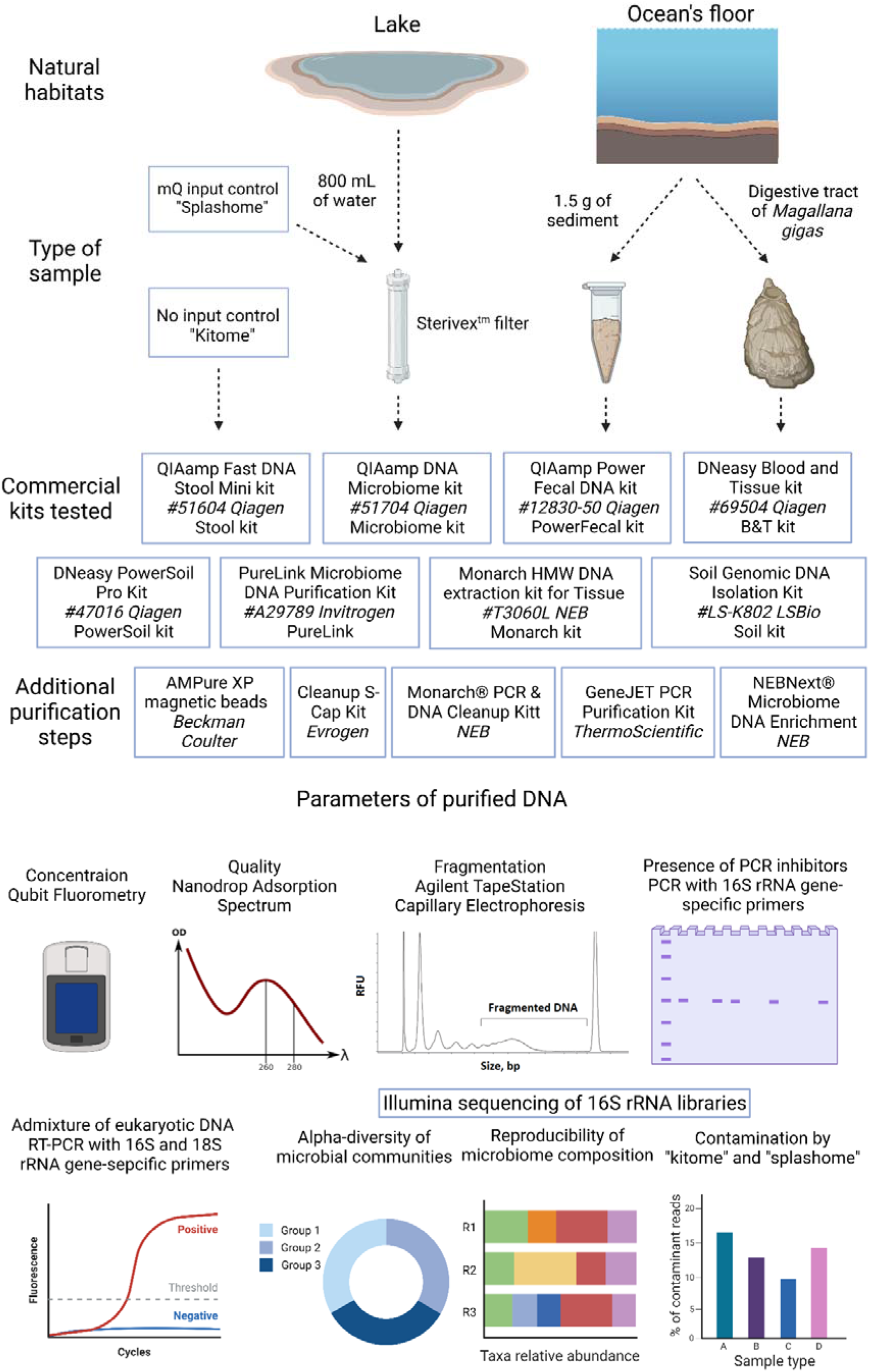
A scheme representing the methodology of the study. Three types of samples were collected. Each sample, along with two types of negative controls (No-input NC; Milli-Q NC), was processed in triplicates with eight commercial microbial DNA purification kits. Additional purification steps were applied to some of the resulting DNA samples to follow the removal of contaminants and reduction of eukaryotic DNA load. All samples were evaluated for an indicated set of parameteres to select the best DNA purification strategy.

In this work, we applied eight commercially available and widely used microbial DNA purification kits to three types of samples: fresh water, deep sea sediments, and digestive system of a model marine invertebrate – Pacific oyster *M. gigas* (Fig. 1). For each sample-kit combination, we estimated: 1) DNA quantity; 2) DNA purity; 3) DNA fragmentation; 4) presence of PCR inhibitors; 5) admixture of eukaryotic DNA; 6) contamination by “kitome” and “splashome; 7) reproducibility of the 16S rRNA amplicon sequencing in 3 biological replicates; 8) alpha-diversity of the sample. The results allowed us to rank each kit and build a comprehensive description matrix that should aid in the selection of the best DNA isolation method for a specific sample type and purpose.

## Materials and Equipment

### DNA purification kits

All DNA purification steps were performed according to manufacturer’s recommendations. When applied, we specifically mention some minor changes introduced into the protocols. General description of each kit and its short identification name used throughout the paper is provided below. When bead beating step was necessary, we used TissueLyser LT (Qiagen) and treated samples at conditions of 50 Hz for 10 minutes, unless other time was specified in the protocol.

**QIAamp Fast DNA Stool Mini kit** (#51604 Qiagen) = **Stool** kit. Purification protocol begins with absorption of the compounds that can degrade DNA and inhibit downstream enzymatic reactions. Next, microbial cells are lysed, and proteins are degraded in the presence of Proteinase K. Then, the sample is loaded onto the QIAamp spin column. DNA bound to the silica membrane is washed two times and concentrated DNA is then eluted.

**QIAamp DNA Microbiome kit** (#51704 Qiagen) = **Microbiome** kit. The first step is lysis of eukaryotic cells and degradation of host nucleic acids with benzonaze. The second step is disruption of bacterial cells through bead beating. Then, the sample is loaded onto the QIAamp UCP mini column. DNA bound to the silica membrane is washed two times and concentrated DNA is then eluted.

**QIAamp PowerFecal DNA kit** (#12830-50 Qiagen) = **PowerFecal** kit. The first step of the protocol is mechano-chemical cell disruption through bead beating in lysis buffer. The Inhibitor Removal Technology is then used to remove common substances that interfere with downstream PCR applications. DNA from supernatant is captured on MB Spin Column, washed two times and eluted from the silica membrane.

**DNeasy Blood and Tissue kit** (#69504 Qiagen) = **B&T** kit. After cell lysis and protein degradation in the presence of Proteinase K, the sample is loaded onto the DNeasy Mini spin column. DNA is then washed two times and eluted from the silica membrane.

**DNeasy PowerSoil Pro Kit** (#47016 Qiagen) = **PowerSoil** kit. The protocol starts with mechano-chemical cell disruption through beat beating in lysis solution. In this kit, Inhibitor Removal Technology is also applied, which should help in removal of organic and inorganic materials such as humic acids, cell debris, and proteins that could interfere with downstream PCR applications. After that, the sample is loaded onto MB Spin Column with silica membrane. DNA is then washed two times and eluted from the silica membrane.

**PureLink Microbiome DNA Purification Kit** (#A29789 Invitrogen) = **PureLink** kit. Multiple protocols are proposed for this kit, we applied soil procedure for all types of samples. In this protocol, the microorganisms are lysed by a combination of heat, chemical, and mechanical disruption, e.g., bead beating is applied. After that, the sample is treated with a cleanup buffer to eliminate PCR inhibitors. The sample is then applied to a PureLink™ spin column. DNA is washed one time and then eluted from the silica membrane.

**Monarch HMW DNA extraction kit for Tissue** (#T3060L NEB) = **Monarch** kit. We skipped the homogenization step, as *M. gigas* samples were uniformly homogenized at preliminary step, common for all kits, while water and soil samples did not require homogenization with pestle. Then, we followed extraction protocol for bacterial samples. All samples were incubated in a STET solution supplied with lysozyme (see below) and heat-treated as recommended by the manufacturer. Next, lysis master mix solution was added to the samples, followed by proteins and RNA removal. In order to preserve the integrity of DNA, it was purified using glass beads, in contrast to silica membrane utilized in all other kits. DNA was gently washed through rotation and eluted from the beads.

**Soil Genomic DNA Isolation Kit** (#LS-K802 LSBio) = **Soil** kit. The sample is homogenized by bead beating and then treated in a special buffer that contains detergent, which serves the purpose of PCR inhibitors removal, such as humic acids, proteins, polysaccharides, and other contaminants. DNA is further bound to a silica spin-column. After one-time washing, DNA is eluted from the silica membrane.

### Sample collection

Ocean sediment samples were hand-collected with a scoop from a bottom sediment horizon of 0–5 cm at the littoral in Kola Bay (Minkino, 69.00203 N, 33.0201 E) in December 2021 (Fig. S1A). Samples were sterile packed, transferred and stored at −20°C until further processing.

Fresh water samples were collected from Skolkovo pond (Moscow, 55.69491 N, 37.35393 E) in June 2022 (Fig. S1A). For each sample, 800 mL of water from the same batch was filtered through the 0.22 μm Sterivex unit (Merck-Millipore) with a peristaltic pump. 800 mL of Milli-Q water was filtered as a negative control in parallel to capture and investigate “splashome”. Filters were subsequently stored at −20°C until processing.

8 individuals of *M. gigas* were collected in Melkovodnaya Bay (Vladivostok, 43.018832 N, 131.885805 E) in July 2022 (Fig. S1A,B). Digestive tracts were isolated and stored at −20°C until processing.

### DNA extraction

At all stages where the use of water was required, we used sterile nuclease-free water (B1500L, NEB).

#### Sediment samples

For DNA extraction, 1.5 g of thawed soil was used per replicate. Studied sediment material was stored in one tube and it was rigorously mixed beforehand to avoid heterogeneity. DNA was isolated in accordance with protocols recommended by manufacturers. For Monarch kit, homogenization step was skipped.

#### Water samples

Sterivex filter membrane was removed from the cover as described ^31^. The membrane was transferred into a sterile and single-use Petri dish with a cell-coated surface facing up and cut into small pieces. Membrane fragments were transferred into the sterile 2-mL microcentrifuge tube. Before further processing, filter fragments were incubated in 200 μL of the lysis buffer STET (50 mM TrisHCl, pH=8.0; 50 mM EDTA; 5% Triton-X100; NaCl – 200 mM; freshly supplied with 10mg/ml lysozyme) at 37°C for 1 h in a heating block with shaking at 600 rpm. After incubation, all liquid was collected for downstream processing in accordance with the manufacturer’s protocols. For samples, which were processed by Monarch kit, the homogenization step was skipped. For samples, which were processed by Microbiome kit, the homogenization step was done before addition of the ATL Buffer (provided in the Microbiome kit).

#### Gut flora samples

Digestive tracts from eight *M. gigas* individuals were thawed, pooled, and then homogenized for 15 min at 50Hz using TissueLyser LT in the Tissue Disruption Tube (QIAamp Fast DNA Tissue Kit (Qiagen)), containing a single stainless-steel bead. Any other tissue homogenization method can be applied as this step. Resultant homogenate was split into 24 aliquots (three replicates for each of eight kits used). DNA was isolated in accordance with the manufacturer’s protocols. For the Monarch kit, the internal homogenization step was skipped.

#### “Kitomes” and “splashomes”

For each kit, two types of negative controls were prepared. For No-input negative control (for Soil and Gut flora samples) no starting material was added, and DNA was extracted solely from the buffers contained in the kit. For Milli-Q negative control (for Water samples), we took laboratory Milli-Q water purified consequently in two steps using the Barnstead Pacific TII (Thermo Scientific) and Simplicity systems (Millipore). 800 mL of Milli-Q was filtered through Sterivex filter units, and DNA was extracted from filter membranes as described for Water samples processing.

### Additional purification procedures

All additional purification was conducted according to the manufacturer’s instructions. An equal volume of samples was taken for each procedure.

**AMPure XP Reagent** (A63880 Beckman Coulter). The Agencourt AMPure XP purification system utilizes solid-phase reversible immobilization (SPRI) paramagnetic bead technology and optimized buffer for high-throughput purification of DNA fragments. Salts, inhibitors, nucleotides, and enzymes are removed using a washing procedure with magnetic separation, resulting in a purified DNA product.

**Cleanup S-Cap** (BC041L Evrogen). DNA fragments bind to the column membrane in the presence of highly concentrated chaotropic salts and optimal pH of binding buffer. Subsequent wash steps allow to get rid of nucleotides, short fragments of nucleic acids, salts, proteins, inhibitors of enzymatic reactions, and other impurities of organic compounds. DNA elution occurs under slightly alkaline conditions in a low salt buffer.

**GeneJET PCR Purification Kit** (#K0702 Thermo Scientific). DNA combined with the binding buffer is added to a purification column. A chaotropic agent in the binding buffer denatures proteins and together with optimal pH promotes DNA binding to the silica membrane in the column. Impurities are removed with a wash step and purified DNA is then eluted from the column with the elution buffer.

**Monarch PCR & DNA Cleanup Kit** (#T1030L NEB). This method also employs a bind-wash-elute workflow. Binding Buffer is used to dilute the samples and ensure they are compatible for loading onto the proprietary silica column under high salt conditions. The Wash Buffer ensures enzymes, detergents and other low-molecular weight reaction components are removed, thereby providing high-purity DNA after elution.

**NEBNext Microbiome DNA Enrichment Kit** (E2612L NEB). Methylated host DNA in a DNA mixture is selectively bound to the mCpG binding domain of human MBD2-Fc protein. After capture, the microbial DNA which is not CpG methylated, or is minimally CpG methylated, remains in the supernatant with minimal sample loss.

**RNaseA** (#T3018-2 NEB). Co-purification of RNA during DNA extraction is a common problem that leads to the overestimation of DNA yield and complication in NGS library preparation. Most commercial kits utilize low alcohol binding conditions, that should result in reduced RNA co-purification. However, an additional RNaseA treatment might be required for samples with obvious saturation in low molecular weight fragments. 10 μL of sample was incubated with 0.4 μL of RNaseA per at room temperature for 10 minutes.

### DNA quantification and quality assessment

DNA concentration was measured using the Qubit dsDNA High Sensitivity Assay Kit or Qubit dsDNA Broad Range Assay Kit on the Qubit 3.0 Fluorometer (Invitrogen). All DNA samples were eluted with 60 μL of elution buffer provided in the corresponding kit so the total amount can be directly compared within one sample type. Detection limit of DNA concentration in the sample for Qubit dsDNA High Sensitivity Assay Kit is 0.1 ng/μl. Thus, samples that were not measurable by this assay should contain less than 6 ng of total DNA. We consider samples with DNA yield above 1500 ng as “passable” for downstream short-read and long-read DNA sequencing.

DNA purity was assessed by measuring the ratios of absorbance at 260 nm to 230 nm (260/230) and 260 nm to 280 nm (260/280) using the NanoDrop1000 spectrophotometer (Thermo Scientific). Samples with a 260/230 ratio between ∼2.0-2.2 and 260/280 ∼1.7-2.0 are assumed as “pure”.

The integrity of genomic DNA was assessed using Agilent TypeStation 4150 (Agilent Technologies) with Genomic DNA ScreenTape System according to the manufacturer’s instructions. 1 μl of sample was used. Samples with DIN above 7.0 were assumed as “high-quality” and acceptable for long-read sequencing.

### PCR Amplification

16S rRNA V3-V4 region was amplified using universal Illumina V3/V4 PCR primers^32^. Each PCR reaction contained 1 μL of DNA, 0.25 μL of forward and reverse primer (final concentration of 0.5 mM), 5 μL of Phusion High-Fidelity PCR Master Mix with HF Buffer (NEB) and 3 μL of nuclease-free water (NEB) for final reaction volume of 10 μL. PCR conditions were 98°C for 30 s, followed by 25 cycles of 98°C for 10 s, 55°C for 20 s, and 72°C for 30 s, with a final extension time of 5 min at 72°C. Amplicons were visualized on a 1% agarose gel containing 0.25 μg/μL ethidium bromide running in 1xTris-EDTA buffer at 100V with a target product size in a range of 400-500bp^32^. Products of successful amplification were clearly visible on gel, otherwise the initial DNA sample was diluted 10 or 100 times for PCR re-examination.

### Quantitative real-time PCR

To estimate a ratio between bacterial DNA and co-extracted eukaryotic DNA, quantitative real-time PCR (qPCR) targeting different variable regions of 16S rRNA and 18S rRNA genes was performed using universal primers (Table 1). Since many DNA samples contained PCR inhibitors, they required additional dilution in order to achieve a robust amplification, yet the same DNA batch was always used for 16S and 18S amplification. qPCR reactions were performed in technical triplicates in optical 96 well plates using the QuantStudio 3 Real-Time PCR System (Applied Biosystems). Each reaction was performed in a final volume of 10 μL, containing 1 μL of template DNA, 5 μL of 2x iTaq Universal SYBR Green Supermix (Bio-Rad), 0.5 μl of forward and reverse primers (0.5 mM final concentration). qPCR protocol consisted of initial denaturation at 95°C for 5 min, followed by 40 cycles of 95°C (30 s), 55°C (20 s), and 72°C (30 s), and the final generation of the dissociation melting curves to verify amplification specificity. The load of eukaryotic DNA was estimated as the difference in the threshold cycle (Ct value) obtained for each sample with 16S- and 18S-specific primers (Ct(18S)–Ct(16S))^33^.

**Table 1.**
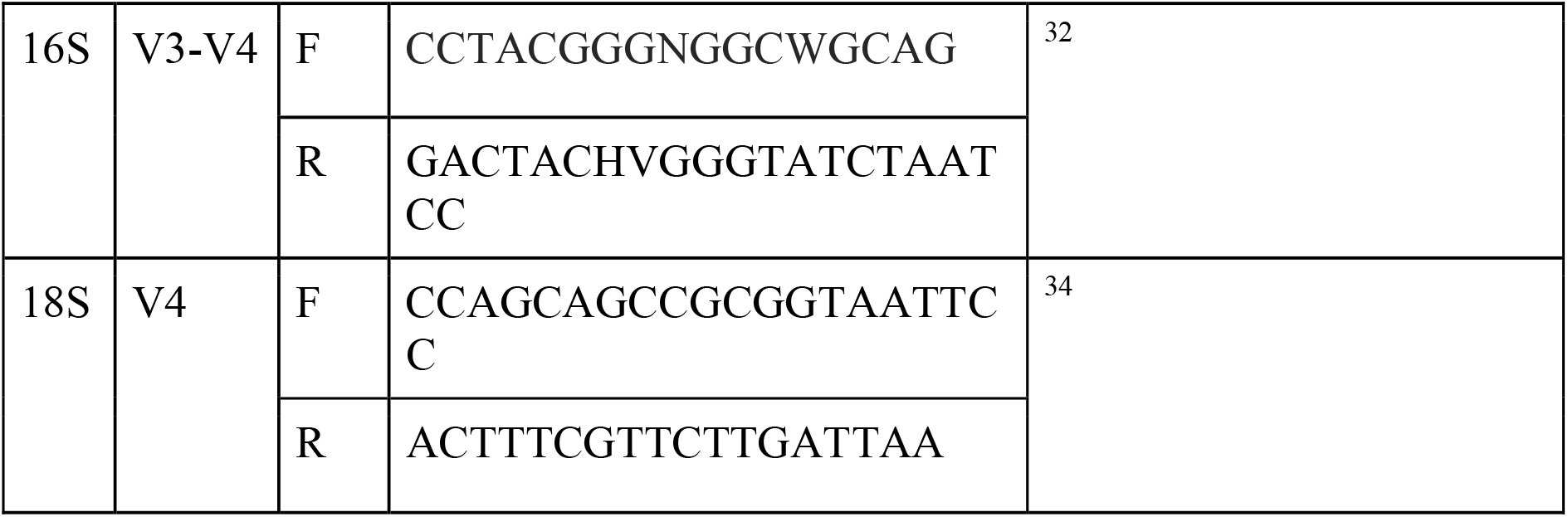
Primers used in the study.

To test the amplification efficiency of the selected 16S and 18S-specific primer pairs, qPCR reactions were performed with five ten-fold dilution series (ranging from 10 to 0.0001 ng/μl) of, correspondingly, *Escherichia coli* and *Homo sapiens* genomic DNA. Primer pairs efficiencies were higher than 98% and were calculated as previously described^33^.

### 16S rRNA libraries sequencing

Amplification of the V3-V4 region of 16S rRNA and library preparations were performed according to the Illumina manual^32^. The amplicon libraries were barcoded, pooled in a single batch and sequenced in Evrogen using NovaSeq 6000, 2 x 250 bp paired-end protocol.

### 16S data analysis

For 16S data analysis, a snakemake pipeline was developed (https://github.com/sutormin94/16S_analysis/). Briefly, quality of sequencing data was checked using FastQC and then reads were trimmed and filtered using Trimmomatic v. 0.39 (SE -phred 33 HEADCROP 17 ILLUMINACLIP:2:30:10 MINLEN:150). Only forward reads were used for downstream analysis. Reads that passed quality control were processed with DADA2 pipeline v. 3.6.2. Resultant amplicon sequence variants (ASVs) were clustered using MMseqs2 v. 10-6d92c (coverage > 0.95, identity > 0.98) and representative sequences were further treated as operative taxonomic units (OTUs). OTUs were returned to DADA2, and taxonomy was assigned to OTUs using the SILVA SSU database v.138^35^. Contamination was removed using R package decontam v. 1.14.0 in the “either” mode with a threshold 0.5^24^. PCoA (Principal coordinates analysis), alpha-diversity, and taxonomic analyses were performed with R packages phyloseq v. 1.30.0^36^, ggplot2 v. 3.3.6 (https://github.com/tidyverse/ggplot2), dplyr v. 1.0.10 (https://github.com/tidyverse/dplyr), vegan v. 2.6.2 (https://github.com/vegandevs/vegan). Shannon index, calculated by phyloseq, was used as a metric that incorporates richness and dominance of OTUs in a community.

### Shotgun sequencing

Libraries for shotgun sequencing were prepared from 300–500 ng of staring DNA using MGIEasy Universal DNA Library Prep Set (MGI Tech), following the manufacturer’s instructions. DNA was sonicated using a Covaris S-220 followed by selection of 200-250 bp-long fragments on magnetic beads (MagBio). The concentration of the prepared libraries was measured using Qubit Flex (Life Technologies) with the dsDNA HS Assay Kit. The quality of the prepared libraries was assessed using Bioanalyzer 2100 with the High Sensitivity DNA kit (Agilent). DNA libraries were further circularized and sequenced by a paired end sequencing using DNBSEQ-G400 with the High-throughput Sequencing Set PE100 following the manufacturer’s instructions (MGI Tech) with an average coverage of 100x. FastQ files were generated using the zebracallV2 software (MGI Tech).

## Results

### Sample collection and processing

The overall scheme of our study is presented in Fig. 1. We processed three types of samples: fresh water, sea sediments, and digestive system of a marine invertebrate (“gut flora”). Given the low titer of prokaryotic cells in the Arctic Ocean water masses, we decided to validate DNA purification from water using a richer sample from a freshwater lake with an intermediate level of algal bloom (Fig. S1A). 800 mL water aliquots collected from a single batch were filtered through Sterivex filters using a peristaltic pump. 800 mL aliquots of Milli-Q water were loaded on separate series of filters as negative controls (“Milli-Q negative controls”) and processed before experimental water samples. All filters were frozen at −20°C and then processed together in a single day. STET lysis solution supplemented with lysozyme was added to fragmented filter membranes and collected liquid served as input for downstream processing (see Methods). The sea sediment sample was collected at the littoral zone of the White Sea (Arctic Ocean water area) (Fig. S1A). The sample was thoroughly mixed to avoid granulometric inhomogeneities, and equivalent of 1.5 grams of dry weight was taken per replicate for processing with each kit. To study the efficiency of microbial DNA purification from marine invertebrates, we selected eight individuals of giant Pacific oyster *M. gigas* (also known as *C. gigas*, NCBI:txid29159) collected at the shores of the Sea of Japan (Pacific Ocean water area) (Fig. S1A,B). The digestive tracts were extracted, homogenized (see Methods) and pooled together for downstream processing (300 mg of homogenate was taken per replicate). All experimental samples were treated in triplicates. Additional negative controls with no input material (“No-input negative controls”), which allowed us to estimate the “kitome” composition, were processed in parallel.

Each sample was purified with eight commercially available kits. Five kits included a beat beating stage for homogenization of the sample (Qiagen’s Microbiome, PowerSoil, PowerFecal, LSBio’s Soil, and Invitrogen’s PureLink) while the other three did not (Qiagen’s B&T and Stool, NEB’s Monarch HMW Tissue). Sample processing was carried out according to manufacturers’ instructions with minor alterations. Purified DNA was eluted in 60 μL of kit elution buffers and stored at −20°C. Samples were evaluated for various DNA parameters and used for 16S rRNA sequencing.

Since sea sediment samples produced good quantities of DNA, yet contained PCR inhibitors (see below), we re-purified these samples using several different protocols (Fig. 1) and repeated our analyses. To assess if the presence of PCR inhibitors affected efficiency of NGS library preparation, we subjected one initial and one re-purified DNA sea sediment sample for shot-gun BGI sequencing. In addition, we evaluated the efficiency of host DNA removal by the NEBNext Microbiome DNA enrichment kit for *M. gigas* samples.

### DNA yield

The concentration of DNA isolated by each kit was determined with Qubit dsDNA High Sensitivity or Broad Range Assay on the Qubit 3.0 Fluorometer. With every sample type, we observed remarkable differences in DNA extraction efficiency between different kits (Fig. 2A, Table 2). For sediment samples, PowerFecal and PowerSoil kits produced the highest yields, while Monarch and PureLink treated samples did not contain detectable quantities of DNA. For water samples, PowerFecal and PowerSoil were also the most efficient and on par with the B&T kit, which is often used for DNA purification from water samples^37, 38^. Notably, in the case of this type of sample, all tested extraction kits isolated measurable quantities of DNA, though Monarch and PureLink continued to be the least efficient. For *M. gigas* samples, only Monarch kit failed to extract DNA. PowerSoil and PowerFecal resulted in high yields, suggesting their potential applicability for different types of samples. The highest amount was isolated with the Stool kit. In this case the amount of isolated DNA exceeded that obtained with other kits more than 10-fold. Since a low molecular weight shmear was evident after gel electrophoreses of Stool kit purified DNA, we checked for RNA contamination by treating the purified DNA with DNase or RNase. The result confirmed that RNA was abundant in the sample (Fig. S2).

**Fig. 2.**
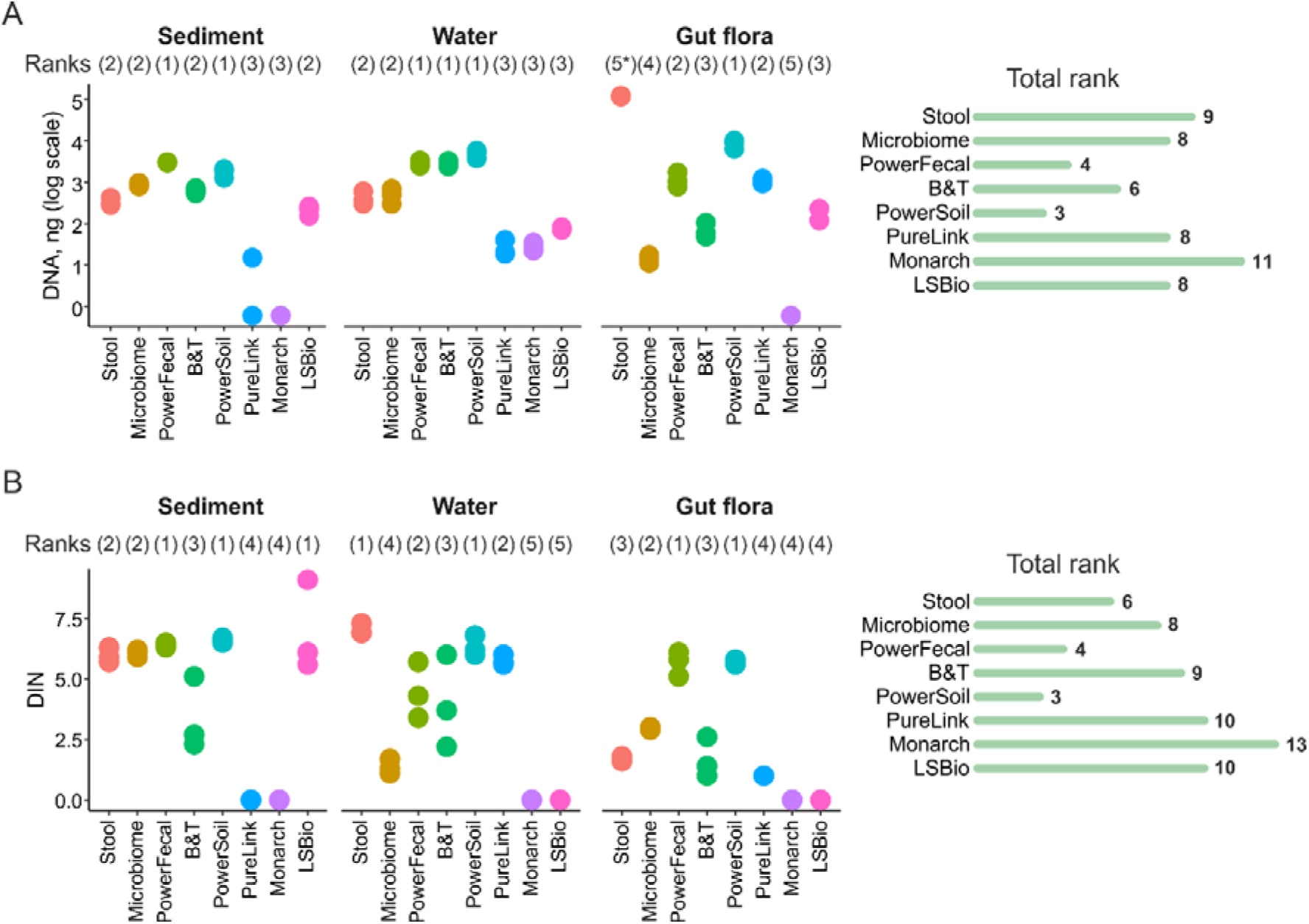
**A)** Total DNA amounts purified by different kits from three types of samples. **B)** DNA integrity numbers (DINs) obtained with three types of samples subjected to purification by different kits. Data for three technical replicates are shown. Right - kit ranking (a sum of ranks for different sample types). Lower ranks indicate higher DNA yeild. (5*) DNA extracted from gut flora samples with a Stool kit was of low quality and was heavily contaminated with RNA.

**Table 2.** Quality and quantity of DNA purified from three types of samples by various DNA extraction kits.

| Kit | Stool | Microbiome | Power Fecal | Blood& Tissue | Power Soil | PureLink | Monarch | Soil |
| --- | --- | --- | --- | --- | --- | --- | --- | --- |
| Sample |  |  |  |  |  |  |  |  |

| Kit<br>Sample |  | Stool | Microbiome | Power Fecal | Blood& Tissue | Power Soil | PureLink | Monarch | Soil |
| --- | --- | --- | --- | --- | --- | --- | --- | --- | --- |
| Sediment | DNA amount, ng | 336.4 ± 75.3 | 858 ± 99 | 3036 ± 67 | 628 ± 90 | 1764 ± 430 | <6 | <6 | 211.2 ± 53.9 |
|  | Quality (260/280) | 1.81 ± 0.01 | 1.38 ± 0.03 | 1.74 ± 0.08 | 1.36 ± 0.02 | 1.89 ± 0.03 | 1.29 ± 0.26 | 1.3 ± 0.6 | 1.62 ± 0.16 |
|  | Quality (260/230) | 1.02 ± 0.10 | 0.69 ± 0.22 | 1.26 ± 0.34 | 0.65 ± 0.01 | 0.22 ± 0.13 | 0.44 ± 0.27 | 0.52 ± 0.07 | 0.38 ± 0.32 |
|  | Fragmentation (DIN) | 5.97 ± 0.31 | 6.07 ± 0.15 | 6.4 ± 0.1 | 3.37 ± 1.51 | 6.6 ± 0.1 | NA | NA | 6.93 ± 1.89 |
|  | 18S<sub>Ct</sub> – 16S<sub>Ct**</sub> | 12.98 ± 1.04 | NA | 9.65 ± 0.49 | NA | 8.45 ± 0.47 | 6.33 ± 1.32 | 7.77 ± 1.02 | 11.94 ± 0.96 |
| Water | DNA amount, ng | 423.2 ± 156.8 | 494.8 ± 196.2 | 2828 ± 485 | 2896 ± 442 | 4960 ± 1042 | 26.4 ± 12.0 | 28 ± 6 | 75.6 ± 5.1 |
|  | Quality (260/280) | 2.03 ± 0.02 | 2.02 ± 0.08 | 2.05 ± 0.33 | 1.88 ± 0.02 | 1.85 ± 0.01 | 2.02 ± 0.73 | 1.7 ± 0.96 | 1.72 ± 0.20 |
|  | Quality (260/230) | 1.63 ± 0.59 | 0.82 ± 0.41 | 1.95 ± 0.22 | 1.69 ± 0.10 | 1.21 ± 0.09 | 0.92 ± 0.67 | 0.08 ± 0.06 | 0.09 ± 0.03 |
|  | Fragmentation (DIN) | 7.03 ± 0.23 | 1.37 ± 0.31 | 4.46 ± 1.16 | 3.97 ± 1.91 | 6.33 ± 0.42 | 5.85 ± 0.21 | NA | NA |
|  | 18S<sub>Ct</sub> – 16S<sub>Ct*</sub> | 5.43 ± 0.42 | 6.37 ± 0.5 | 4.47 ± 0.34 | 3.53 ± 0.29 | 5.2 ± 0.29 | 7.37 ± 0.46 | 3.58 ± 1.51 | 6.77 ± 1.37 |
| *M. gigas* | DNA amount, ng | 119760 ± 4380 | 14.3 ± 3.1 | 1196 ± 501 | 71.2 ± 30.0 | 1038 ± 105 | 1096 ± 151 | <6 | 156.8 ± 63.7 |
|  | Quality (260/280) | 0.05 ± 0.04 | 1.83 ± 0.11 | 1.71 ± 0.01 | 1.50 ± 0.03 | 1.94 ± 0.06 | 1.80 ± 0.03 | 1.17 ± 0.41 | 1.91 ± 0.13 |
|  | Quality (260/230) | 0.05 ± 0.04 | 0.13 ± 0.06 | 0.86 ± 0.18 | 0.51 ± 0.06 | 1.12 ± 0.67 | 1.68 ± 0.06 | 0.67 ± 0.21 | 1.07 ± 0.81 |
|  | Fragmentation (DIN) | 1.7 ± 0.1 | 2.93 ± 0.06 | 5.67 ± 0.51 | 1.67 ± 0.83 | 5.73 ± 0.12 | 1.0 ± 0.0 | NA | 1.0 ± 0.0 |
|  | 18S<sub>Ct</sub> – 16S<sub>Ct**</sub> | -8.2 ± 0.16 | 4.56 ± 1.31 | -7.14 ± 0.38 | 6.31 ± 1.12 | -8.47 ± 0.41 | 0.83 ± 1.3 | 3.76 ± 1.23 | 2.88 ± 1.67 |

\* - 10x dilution was taken for qPCR reaction due to the presence of PCR inhibitors.
\*\* - 100x dilution was taken for qPCR reaction due to the presence of PCR inhibitors.
NA – No result was obtained due to low DNA quality or the presence of PCR inhibitors.

### DNA fragmentation

The extent of DNA fragmentation was estimated by capillary electrophoresis on the TapeStation 4150 System (Agilent). This analysis allows one to calculate the DNA integrity number (DIN). DINs above 7 (roughly corresponding to samples with median fragment lengths above 15 kbp) are usually considered applicable for long-read (ONT or PacBio) sequencing^39, 40^. For complex environmental samples, such quality of DNA is rarely achieved, and storage of frozen samples can result in increased fragmentation. Extensive DNA fragmentation (DIN below 3) can affect all types of downstream applications: long-read, short-read, and even 16S rRNA sequencing^40–42^. We measured DIN after purified DNA samples were kept frozen at −20°C for 3 months (Fig. 2B, S3 and Table 2). For sediment samples, all kits that extracted measurable quantities of DNA, except B&T kit, demonstrated acceptable DINs of ∼6-7. Stool and PowerSoil kits resulted in DINs of ∼6-7 for water samples, while the commonly used B&T kit produced DNA with a DIN of ∼4. *M. gigas* samples demonstrated intense fragmentation, reflecting possible contamination with host-derived nucleases. The highest DIN for these samples (∼5.5) was achieved by PowerFecal and PowerSoil kits. Unexpectedly, the Monarch HMW kit, designed for purification of high molecular weight DNA from tissue samples, performed poorly resulting in DNA quantities that could not be measured on the TapeStation (Fig. 2).

### DNA purity and presence of PCR inhibitors

The purity of DNA samples was measured on a NanoDrop spectrophotometer. The 260/280 absorption ratio of 1.8-1.9 is usually considered as evidence of absence of protein contamination, while the 260/230 absorption ratio in a range of 1.8-2.5 demonstrates the lack of organic impurities^43^. In general, all purified DNA samples lacked protein contamination but contained excess compounds absorbing at 230 nm (Fig. S4). Stool, PowerFecal and PowerSoil kits performed best for sediment samples; Stool, PowerFecal and B&T– for water samples; while PureLink, PowerFecal, and PowerSoil produced best results with *M. gigas* samples.

As inhibition of downstream enzymatic reactions is the major adverse effect of DNA contaminants, another way to assess DNA purity is to estimate the efficiency of PCR amplification with purified DNA samples. To do so, we performed PCR using universal 16S rRNA gene-specific primers and 25 amplification cycles with Phusion DNA polymerase (NEB) according to manufacturer’s recommendations. For all sediment and *M. gigas* samples, except for those purified with PowerSoil, we failed to identify the expected amplicon, indicating the presence of PCR-inhibiting impurities (Fig. 3, S5, Table 2). Although water samples were expected to contain less PCR inhibitors, samples purified with Stool, Microbiome and B&T kits also failed to produce a PCR product. A common way to overcome PCR inhibition issue for DNA obtained from natural sources is dilution of samples, which lowers concentration of inhibitors below a threshold while still providing sufficient DNA for amplification^44, 45^. Upon 100x dilution, all samples (except the negative “kitome” controls) produced 16S rRNA gene amplicons. Thus, we used a dilution factor as a rough estimate for the presence of PCR-inhibiting impurities (Fig. 3, S5, Table 2). PowerSoil, PowerFecal and Stool kits include a specific inhibitors removal step, and indeed PowerSoil performed best in this comparison.

**Fig. 3.**
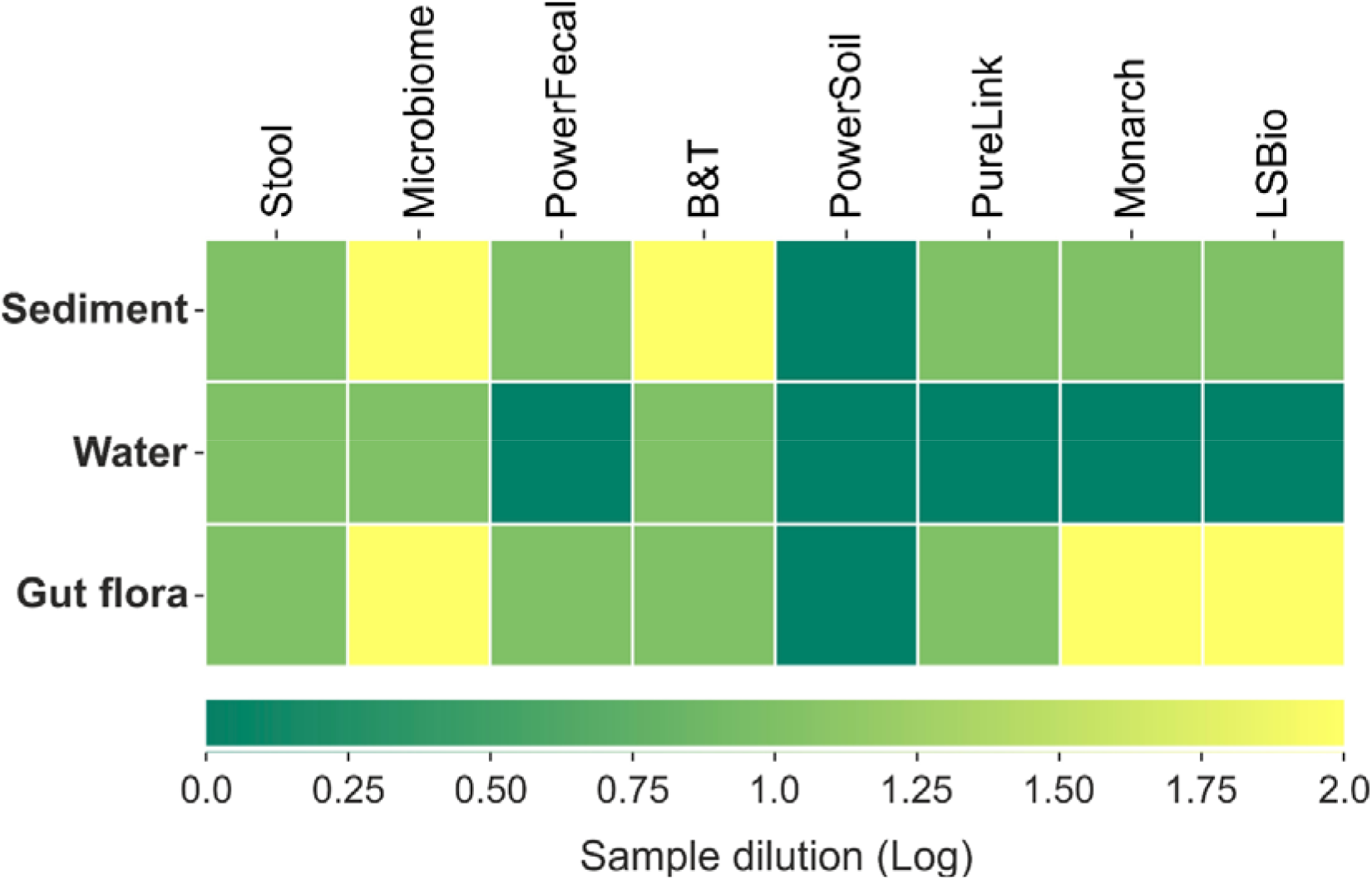
The presence of DNA inhibitors estimated as a dilution factor required to achieve visible production of a 16S rRNA amplicon. Mean results obtained for the three technical replicates are shown.

Since contamination with PCR inhibitors represented a serious issue for the sea sediment samples, we investigated if additional purification steps could help to cope with this problem. We selected DNA samples purified with Stool and Microbiome kits, which required 10x or 100x dilution for efficient 16S PCR amplification, respectively. Samples were divided in equal aliquots and re-purified using protocols described in the Methods section. For each re-purification, 50 ng (Stool) or 150 ng (Microbiome) of DNA was taken and retention efficiency was close to 100 % (Fig. S5A). To access efficiency of PCR inhibitors removal, we performed 16S PCR with non-diluted and 10x diluted non-purified and re-purified samples. Both input samples did not produce the expected PCR product (Fig. S5B). In the case of Microbiome samples, only columns re-purification with Evrogen kit led to the appearance of 16S gene amplicons, while for Stool kit-purified samples all additional purifications protocols used were efficient (Fig. S5C).

We also investigated whether the presence of PCR inhibitors affects the quality of DNA libraries for shotgun sequencing on the BGI platform. A DNA sample purified from sediments with the PowerFecal kit (which required a 10x dilution for efficient PCR) and a corresponding sample re-purified with DNA Purification kit (Evrogen) (no dilutions required for efficient PCR) were used. Both samples were prepared according to a standard procedure that involved AMPure XP beads purification (see Methods). Samples passed through library preparation steps with similar efficiency and produced assemblies of a comparable quality (Fig. S6), suggesting that samples with moderate contamination with PCR inhibitors can be subjected to shotgun sequencing with a standard library preparation method without the additional purification step.

### Admixture of eukaryotic DNA

The quantities of eukaryotic DNA in samples were assessed using qPCR with primers specific for 16S rRNA and 18S rRNA genes. The universal 18S rRNA primers used annealed to the 18S rRNA gene of *M. gigas* genome^46^. The difference in 18S to 16S amplicons abundance was estimated by subtracting threshold cycles (the Ct(18S)–Ct(16S) value). Values above 0 indicate the predominance of 16S rRNA-containing DNA molecules in the sample and 3 Ct cycles are expected to roughly correspond to a 10-fold difference of template molecules. As can be seen from Fig. 4 and Table 2, sediment and water samples produced a ∼100-1000x higher 16S rRNA signal, although, given the substantially larger sizes of eukaryotic genomes, this does not directly indicate a similar excess of prokaryotic DNA. For the *M. gigas* samples, many kits demonstrated Ct(18S)–Ct(16S) values above 0, and the best result was achieved with the Microbiome kit which is specifically designed to deplete host DNA. At the same time, Stool, PowerSoil, and PowerFecal kits demonstrated the highest levels of eukaryotic DNA admixture. It should be noted that though the PowerFecal and PowerSoil kits performed best in other tests and produced the largest amount of DNA from *M. gigas* samples. However, the high DNA yields may be due to eukaryotic DNA admixture, suggesting that the weakness of these kits lies in the inability to discriminate between microbial and host DNA. In addition, PureLink-purified *M. gigas* sample showed a Ct(18S)– Ct(16S) value close to 0, and the amount of DNA purified with this kit was significantly higher compared to other kits with intermediate level of eukaryotic DNA admixture. Thus, considering the amount of total DNA and abundance of 16S rRNA in samples, B&T, PureLink, and Microbiome kits performed best for *M. gigas* samples.

**Fig. 4.**
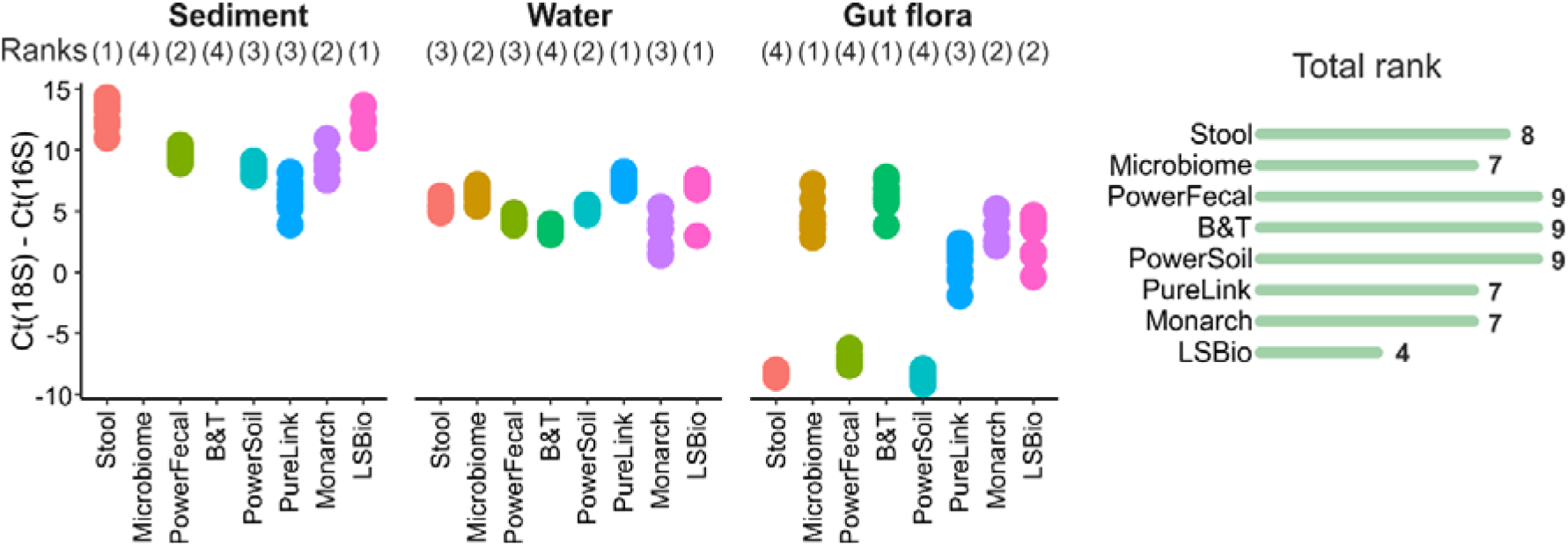
Eukaryotic DNA admixture estimated by the qPCR Ct values obtained with 18S and 16S rRNA gene-specific primers. Data for three technical qPCR replicates for each of the three kit purification replicates are shown. Rigth - total kit rankings (a sum of ranks for different sample types). Lower ranks indicate higher proportion of microbial DNA.

To evaluate whether eukaryotic DNA admixture can be lowered by an additional treatment, we employed the NEBNext Microbiome DNA Enrichment kit, which selectively captures CpG methylated eukaryotic DNA, and re-purified Stool, PowerFecal and PowerSoil samples that contained the highest amounts of eukaryotic DNA. Even though mollusk’s DNA is considered hypomethylated^47^, the treatment of selected samples improved the Ct(18S)– Ct(16S) value by 3 Ct units on average, though microbial DNA still remained significantly underrepresented (Fig. S7).

### Contamination levels and determination of “kitomes”

To further explore technical biases associated with the benchmarked DNA extraction kits, V3-V4 regions of 16S rRNA gene were PCR amplified from DNA extracted from environmental microbial communities and from mock samples prepared without input material (No-input negative controls representing “kitome”) or with sterile Milli-Q water (Milli-Q negative controls representing “kitome” + “splashome”). Obtained 16S amplicons were subjected to HTS using Illumina NovaSeq 6000 and the microbial diversity was studied at the operational taxonomic unit (OTU) level.

For environmental samples, on average, 93054 reads (standard deviation (SD) of 34626 reads) were obtained. For mock samples, as expected, the average number of reads was lower -77564 (SD of 63798 reads). For further analysis, samples with a large number of reads were subsampled to 100,000 reads. Inspection of rarefaction curves at the level of OTUs demonstrated that for all samples, the sequencing depth reached the saturation level (Fig. S8).

In the mock samples, the most abundant taxa were represented by Beta-proteobacteria (*Burkholderia, Ralstonia*), Gamma-proteobacteria (*Acinetobacter*, *Escherichia*/*Shigella*), Actinomycetia (*Cutibacterium*), and other bacterial genera known to be kit and laboratory contaminants (Fig. 5A)^48–51^. A full list of OTUs found in the mock samples can be found in Table S1.

**Fig. 5.**
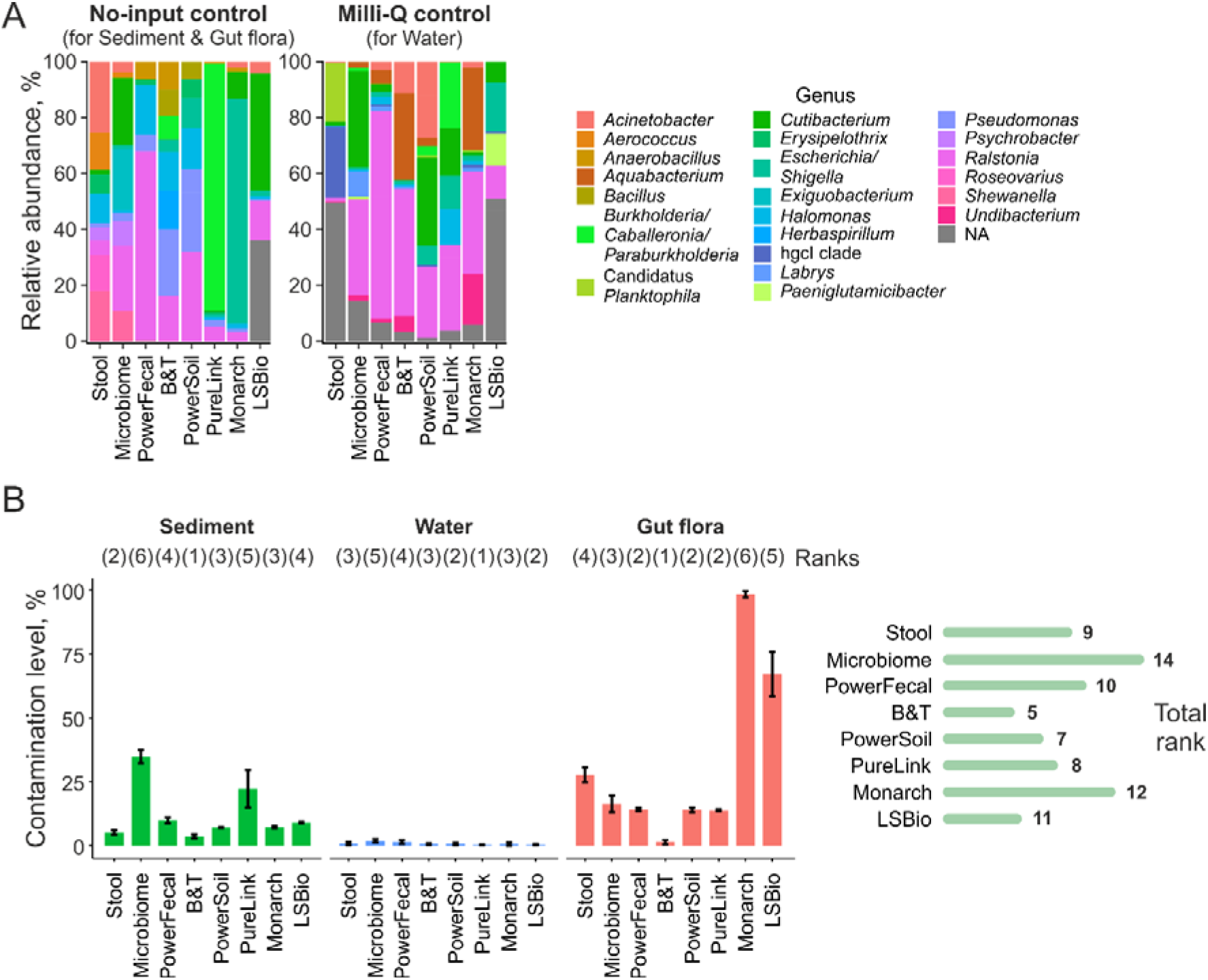
Determination of kitomes of commercial DNA extraction kits. **A**) Diversity of kitomes on a genus level. Data is shown for genera with relative abundances >5%. **B**) Left - contamination levels of natural samples processed with commercial kits. Means of three technical replicates +/- SD are shown. Kit rankings for each sample type are shown above the bar plot. Right - total kit ranking (a sum of ranks for different sample types). Lower ranks indicate lower contamination levels.

Using the data obtained for mock samples, contamination was removed from natural samples with the Decontam package^24^. Decontamination was performed in the “either” mode which utilizes both DNA concentration and the OTU frequency data. In order to effectively remove contamination, the probability threshold was set to an aggressive 0.5 value (compared to 0.1 default value). With this threshold, sequences that are more prevalent in mock rather than in natural samples, are considered as contaminants. The fraction of OTUs classified as contamination (the sum of contaminant OTU frequencies divided by the total frequency in the sample) was calculated for environmental samples and is further referred to as “contamination level”. Contamination levels varied significantly for different sample types with water samples associated with lower average level of contamination (0.8% vs 12% for sediment and 32% for gut flora samples), indicating that some sample types may be more prone for contamination than others. The increased contamination levels in samples from digestive tract of *M. gigas* are likely due to low amounts of bacterial DNA in these samples (Table 2). Contamination levels reached 20-80% in sediment samples processed with Microbiome and PureLink kits and in gut flora samples processed with Stool (also reflecting poor quality of this sample, see Fig. S2), Monarch, and LSBio kits (Fig. 5B, S9). After decontamination, the composition of these bacterial communities changed drastically (Fig. S10). The B&T kit demonstrated the lowest level of contamination for sediments and gut flora, while PureLink showed the lowest contamination level for water samples. DNA isolation with B&T and PowerSoil kits was associated with the overall lower contamination levels, while extraction with Microbiome and Monarch kits led to higher contamination levels (Fig. 5B).

### Technical reproducibility of DNA extraction kits

To inspect the effect of DNA extraction kits on the technical variability of bacterial communities’ composition, lists of OTUs were compared within sets of three technical replicates after the decontamination step. Reproducibility level was measured as a ratio of OTUs present in all three technical replicas to the total number of unique OTUs found in at least one replicate. Reproducibility levels varied significantly between sample types with the lowest and the highest levels associated with, correspondingly, gut flora and water samples (Fig. 6A-C, S11). Lower reproducibility of gut flora and sediment samples may be caused by the low amounts of extracted microbial DNA subjected for sequencing (*M. gigas* samples are enriched with host DNA, Table 2) and/or by innate heterogeneity and high diversity of microbial communities (sediment samples, see below). Within sample types, kits demonstrated high variability. For sediment samples, the LSBio and PowerSoil kits designed to extract DNA from soil were ranked the first and the second best, while the PureLink kit performed the worst. For water samples, several kits (Stool, PowerSoil, and Monarch) performed equally well, while PowerFecal and LSBio kits demonstrated the lowest reproducibility levels. For gut flora samples, the PowerFecal and PowerSoil kits shared the first place, while the Monarch kit was the least reproducible, likely reflecting the poor quality of DNA (Table 2). Overall, in terms of reproducibility, PowerSoil outperformed other kits, ranked the first for two out of three sample types, while Microbiome and PureLink kits performed poorly (Fig. 6A).

**Fig. 6.**
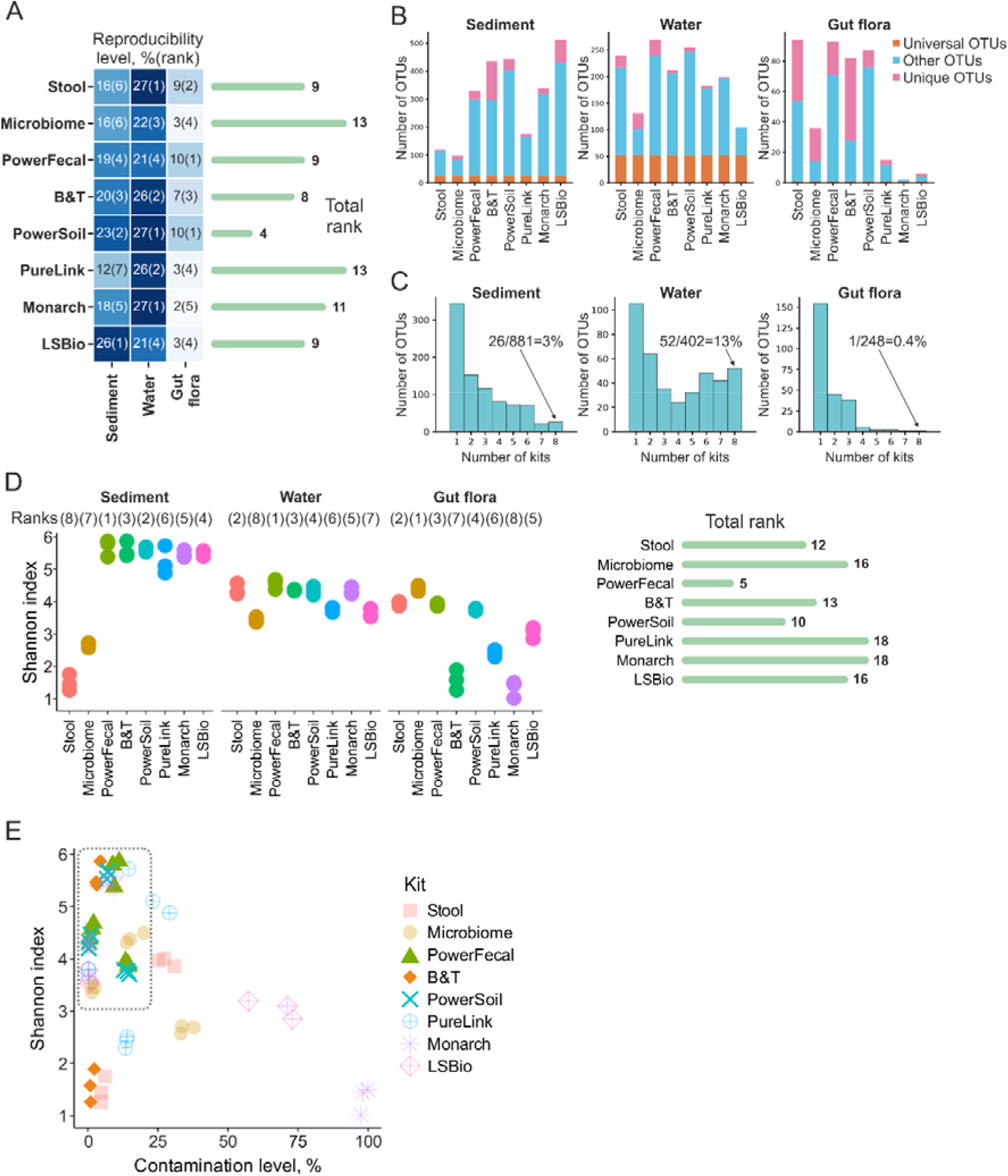
Reproducibility of DNA extraction kits and the effect of kits on alpha-diversity of bacterial communities. **A**) Reproducibility levels of commercial DNA extraction kits for different sample types. Right - total kit rankings (a sum of ranks for different sample types). Lower rankings indicate higher reproducibility levels. **B**) Bar plots representing fractions of OTUs shared between all DNA extraction kits benchmarked for a particular sample type (Universal OTUs, orange), OTUs found by just one kit (Unique OTUs, pink), and other OTUs (found in DNA prepared by two to seven kits, light blue). **C**) Fractions of OTUs shared between different numbers of DNA extraction kits. Numbers of OTUs found by all 8 studied DNA extraction kits are shown. **D**) Microbial alpha diversity (the Shannon index) of DNA samples prepared using different kits. Data for three technical replicates are shown. Right - total kit rankings (a sum of ranks for different sample types). Lower rankings indicate higher Shannon index values. **E**) Contamination level and alpha diversity (Shannon index) of samples processed with different DNA extraction kits. A group of samples with high diversity and low contamination levels is marked with a dashed rectangle.

### Alpha diversity of microbial communities

To assess the effects of DNA extraction kits on the diversity of microbial communities, alpha diversity (the Shannon index) was calculated for purified DNA samples after the contamination removal step (Fig. 6D). Higher Shannon indexes were observed for sediment samples and lower Shannon indexes were observed for gut flora samples, indicating high and low complexities of the two communities, correspondingly. The Shannon index was significantly decreased for samples processed with Microbiome (sediment and water samples), PureLink and LSBio (water and gut flora samples), Stool (sediment samples), and B&T and Monarch (gut flora samples) kits. The Microbiome kit was associated with the highest Shannon index for gut flora samples and the PowerFecal kit was associated with the highest Shannon indexes for soil and water samples. Considering contamination levels and alpha diversities associated with different DNA extraction kits, the PowerSoil and PowerFecal kits resulted in high alpha diversity and low contamination levels for all types of samples analyzed (Fig. 6E).

### Effects of DNA extraction kits on the composition of bacterial communities

To investigate the effects of DNA extraction kits on the composition of microbial communities, beta diversity of natural microbiomes after the decontamination step was assessed using the Bray-Curtis dissimilarity and NMDS for visualization. Visual inspection of relative abundance and NMDS plots (Fig. 7) revealed outlier kits with distorted microbial community composition. All sample types processed with the Microbiome kit, sediment samples processed with the Stool kit, and water samples processed with the LSBio kit had dramatically distorted composition. Moreover, gut flora samples processed with LSBio and Monarch kits had distorted and highly heterogenous compositions, which corresponds to low level of technical reproducibility and poor quality of DNA purified with these kits (Fig. 6A). Using the PERMANOVA method, kit selection was found to be a factor significantly affecting the composition of microbial communities for all sample types (p-values=0.001).

**Fig. 7.**
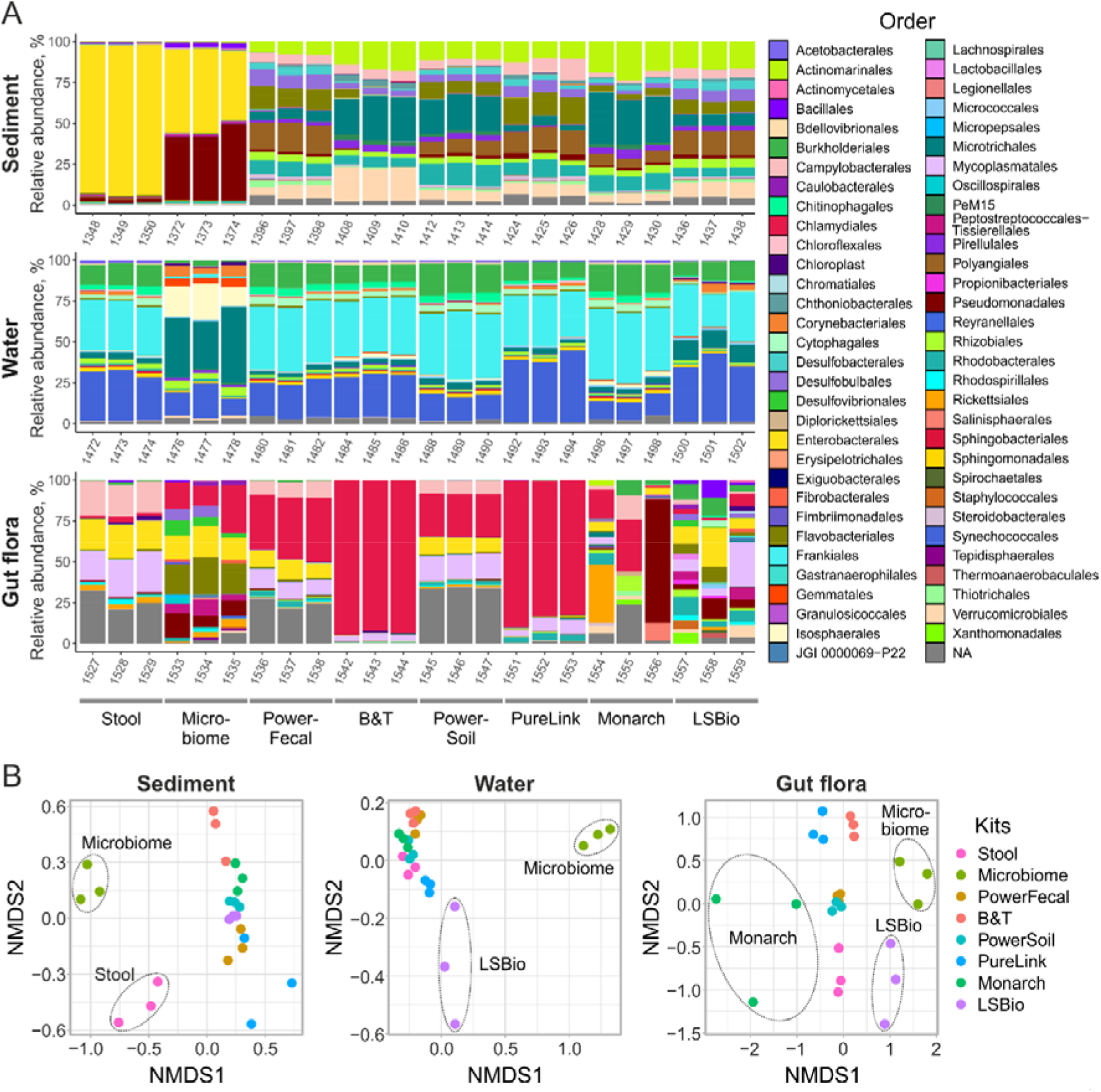
Effects of DNA extraction kits on the composition of bacterial communities. **A**) Microbial communities’ composition of environmental samples at the order level after the decontamination step. Data is shown for all technical replicates independently. Orders with relative abundances >1% are shown. **B**) NMDS plots with points representing microbial communities. Dashed ellipses indicate sample groups distant from the majority of samples.

We further estimated how the communities’ composition overlaps between samples purified by different kits. For this analysis, only OTUs found in 3 replicates of DNA prepared with each kit were considered. As DNA from water samples was efficiently purified by most kits, more than 50 OTUs were identified in all samples, while the level of kit-specific OTUs was low (Fig. 6B, C). For sediment samples, the number of universal OTUs was lower, and the B&T kit purified more unique OTUs than other kits (Fig. 6B, C). Because of the overall lower quality of DNA from *M. gigas* and the failure of some kits to generate detectable amounts of DNA, these samples were the least consistent (Fig. 6B, C). With PureLink, Monarch, LSBio, and Microbiome kits, less OTUs were identified, while increased amount of unique OTUs was observed in Stool and B&T purified DNA.

## Discussion

The results of any metagenomic analysis depend on the quality of input DNA. Given that DNA purification steps introduce multiple biases^52–56^, it is highly advantageous to know the strong sides and limitations of available DNA isolation tools. Comprehensive study of marine microbial communities requires processing of different sample types, including water, sea floor sediments, and symbionts of multicellular organisms. DNA isolation from various types of sample may face challenges, such as low titers of microbial cells, the presence of PCR inhibitors, or eukaryotic DNA. Despite multiple studies discussing the choice of DNA isolation methods for human microbiota^56–59^ and some studies focused on specific marine communities^60, 61^, a holistic investigation of DNA purification kits performance with marine samples is lacking.

To cover this gap, we selected eight commercially available DNA purification kits, and benchmarked them using three sample types (sea sediment, water, and gut flora of *M. gigas*). For each kit-sample combination, we measured the quality of purified DNA and its applicability for 16S metagenomics and described the resulting community composition. We estimated DNA concentration, DNA integrity (DIN), DNA purity (260/230 and 260/280 absorbance ratios), presence of PCR inhibitors, admixture of eukaryotic DNA (the 18S/16S ratio obtained using the Ct(18S)-Ct(16S) value), and three parameters describing the composition of microbial communities revealed by 16S metagenomics: contamination level, alpha diversity of microbial communities, and reproducibility of results. Using these parameters, the kits were ranked according to their performance. Higher DNA yield, DIN, reproducibility level, alpha diversity, and lower presence of inhibitors, 18S/16S ratio, and contamination level were considered to be linked to better performance.

Based on average rankings obtained for three sample types, the PowerSoil and PowerFecal kits, and the Microbiome and Monarch kits had the highest and the lowest performance levels, respectively (Fig. 8, S12). The PowerSoil and PowerFecal kits had the worst 18S/16S ratios, indicating that they may have a bias toward eukaryotic DNA extraction, which, however, did not have apparent negative effects on the profiling of microbial communities.

**Fig. 8.**
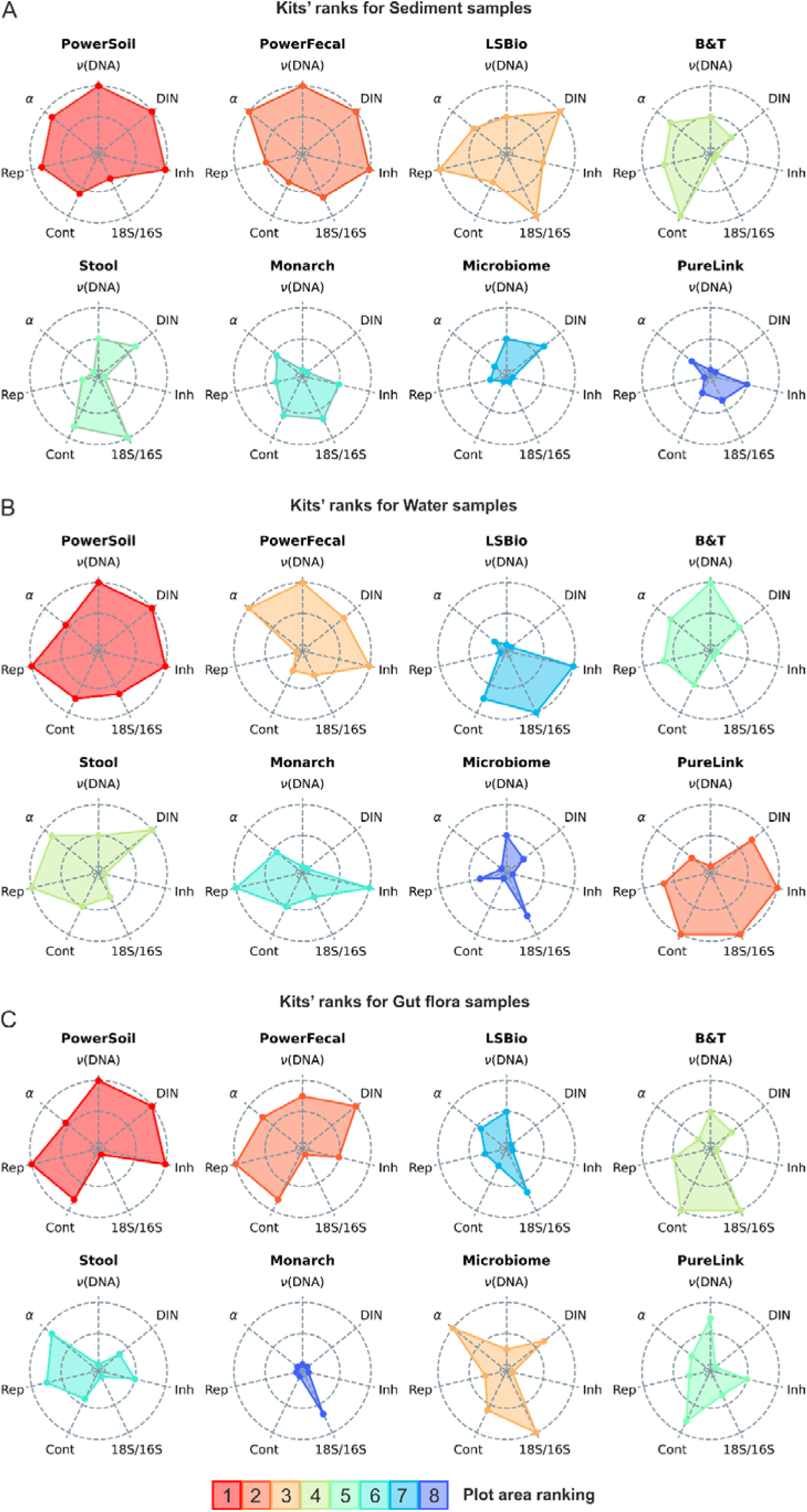
Radar-plots demonstrating the performance of DNA extraction kits with Sediment (**A**), Water (**B**), and *M. gigas* gut flora samples (**C**). L(DNA) – DNA yield, DIN – DNA integrity, Inh – presence of PCR inhibitors (higher rank indicates the lower level of inhibitors), 18S/16S – 18S/16S ratio (higher rank indicates the lower ratio), Cont – contamination level (higher rank indicates the lower level of contamination), Rep – reproducibility level, L – alpha-diversity. Kits were ordered by the sum of rankings.

PowerSoil was ranked the best for all sample types, followed by PowerFecal for soil and gut flora samples and PureLink for water samples (Fig. 8). The LSBio kit designed for DNA extraction from soil was ranked the third for this type of sample (Fig. 8A). The Microbiome kit designed to extract microbial DNA from samples with high eukaryotic load demonstrated the best 18S/16S ratio and high alpha diversity. However, the high level of PCR inhibitors and low reproducibility in our opinion limits its applicability, although by the sum of parameters the kit was ranked third for the gut flora samples (Fig. 8C). Overall, the PowerSoil and PowerFecal kits can be considered as the most versatile and useful for a wide range of environmental samples. The LSBio and PureLink kits can be recommended for DNA extraction from, respectively, soil and water samples.

An inevitable source of biases associated with any DNA purification procedure is the presence of microbial DNA contaminants in kit solutions and columns, which may result in identification of OTUs that are actually absent from the studied sample. The set of such contaminants is referred to as the “kitome”. Using samples with no input material, we describe OTUs specifically associated with each tested kit (Table S1). The list of detected OTUs can be used for decontamination at the stage of 16S data analysis. In addition, we studied the contribution of contamination from the laboratory Milli-Q water – another important source of non-specific 16S signal. Using Milli-Q water as an input, we described the combined composition of the “kitome” (kit-specific) and “splashome” (laboratory-specific) contaminations. The results showed that samples of poor quality in general demonstrated much higher levels of contamination.

Our study has obvious limitations, as a multitude of additional parameters, such as details of sampling, storage and homogenization procedures can affect the quality of purified DNA and communities’ composition ^14, 50, 58, 62, 63^. Also, the usage of environmental samples for kit benchmarking does not allow to identify kit biases for specific bacterial taxa (“taxa-specific biases”), as the “ground truth” composition of investigated communities’ was not known. To overcome this limitation, mock communities with an *a priori* known composition are typically used ^64–66^. Mock communities, however, do not reflect biases associated with natural samples (“sample-specific biases”), e.g., heterogeneity, presence of PCR inhibitors, presence of eukaryotic cells, high complexity, etc. To account for both taxa-specific and sample-specific biases a combined approach is needed, based on benchmarking on mock and environmental communities. Nevertheless, purification of DNA from initially identical replicates of environmental samples with a set of eight DNA purification procedures allowed us to characterize sample-specific biases and highlight kits with the highest performance. While more elaborate DNA purification protocols can be further developed based on standard recommendations for these kits our results should aid in selection of the DNA isolation techniques most appropriate for different types of marine samples and downstream applications (amplicon and shotgun short-read or long-read NGS sequencing).

## Author Contributions

Conceptualization, A.I., D.Sl., and D.Su.; methodology, A.I., A.D., D.Sl., and D.Su.; sample collection, K.O., Y.D., and R.V.; investigation, A.D., D.Sl., V.M., and D.Su.; BGI sequencing, V.B., and D.K.; writing—original draft preparation, A.D., D.Sl., A.I., and D.Su.; writing—review and editing, A.I., D.Su., and K.S.; All authors have read and agreed to the published version of the manuscript.

## Data Availability Statement

Raw and processed 16S rRNA amplicon and shotgun sequencing data reported in the paper has been deposited to the NCBI repository under bioproject ID PRJNA996732.

---

**Fig. S1.**
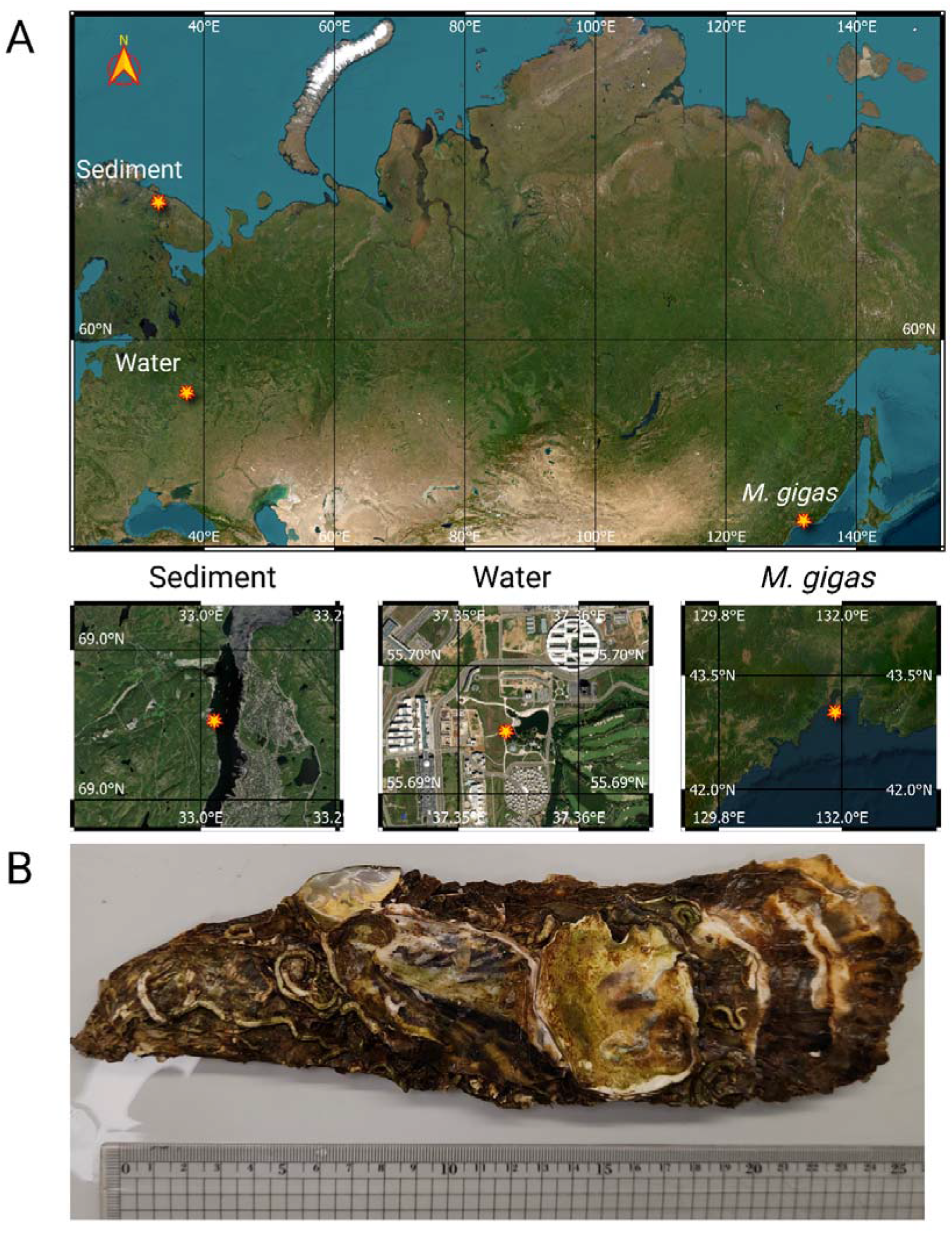
**A)** Geographical locations for water, sea sediment, and Pacific oyster (*M. gigas*) collection spots. Map was prepared using Open Source Geospatial Foundation Project (http://qgis.org) **B)** Selected individual of Pacific oyster (*M. gigas*) aligned with a centimeter ruler.

**Fig. S2.**
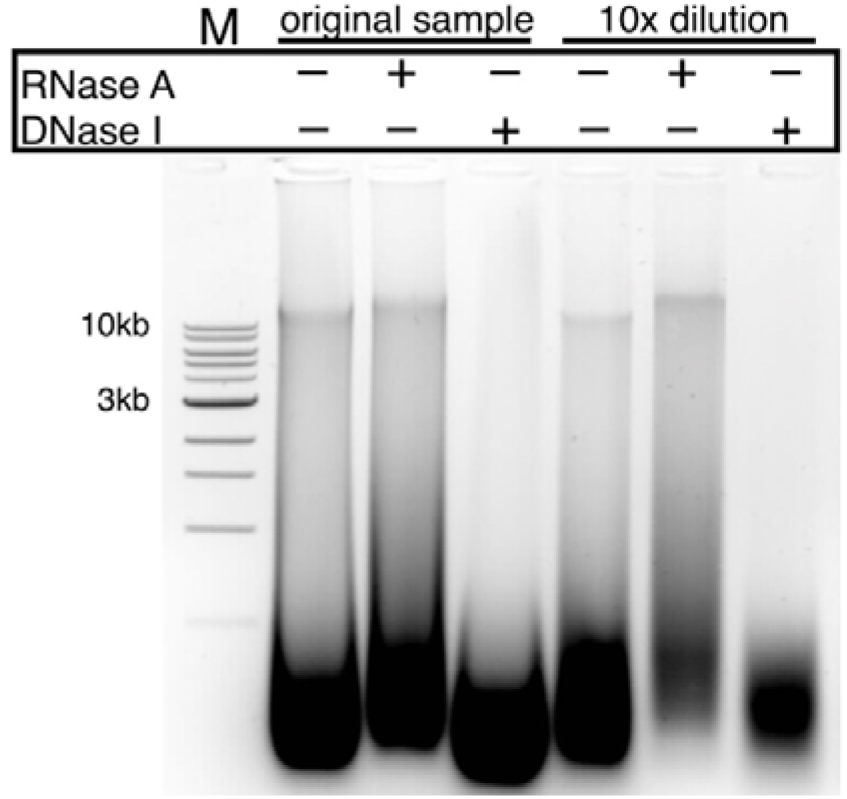
DNase and RNase treatment of *M.gigas* DNA sample isolated with Stool kit.

**Fig. S3.**
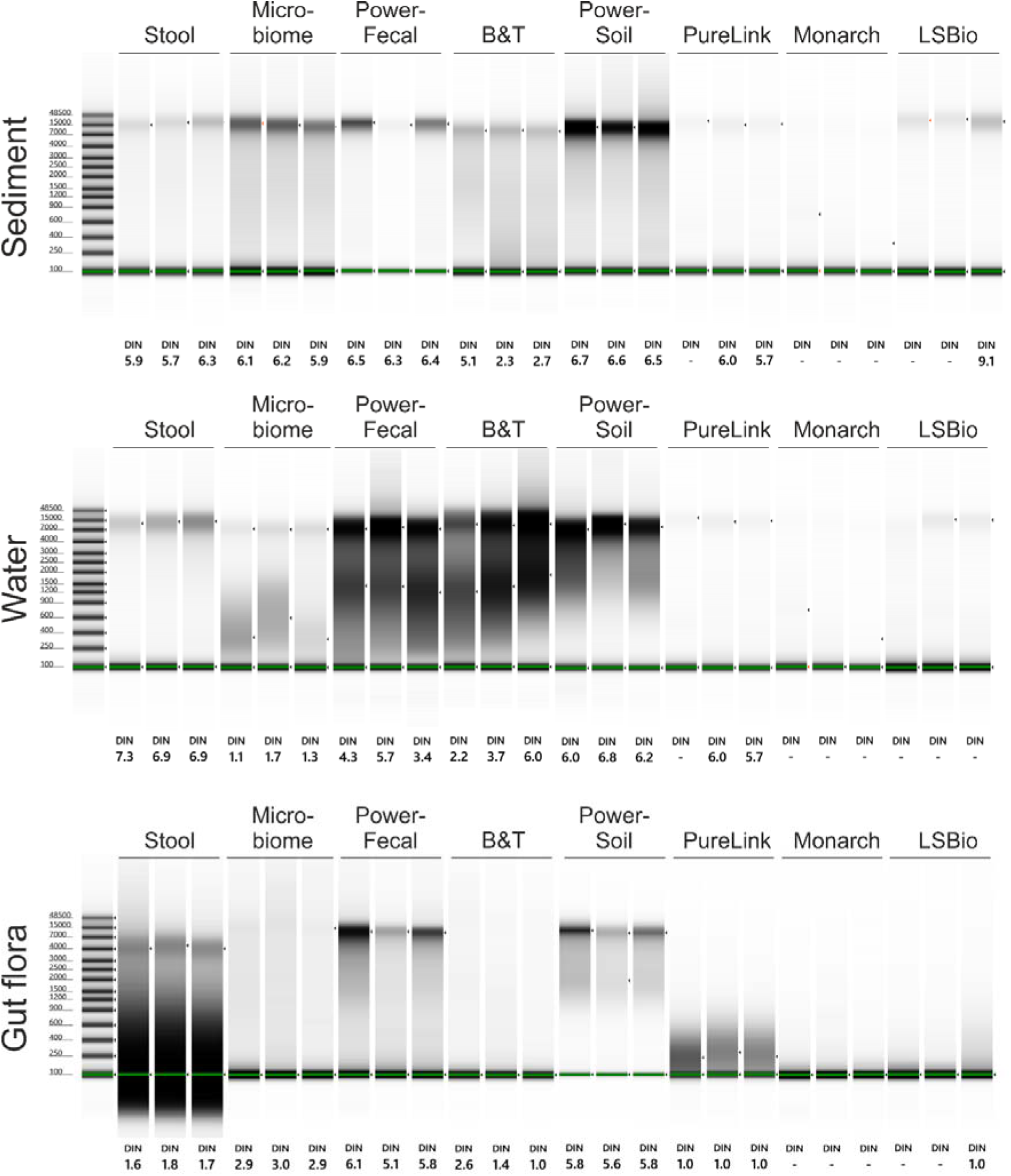
Capillary electrophoresis estimation of the DIN values performed on a TapeStation 4150 (Agilent) with Genomic DNA ScreenTape System.

**Fig. S4.**
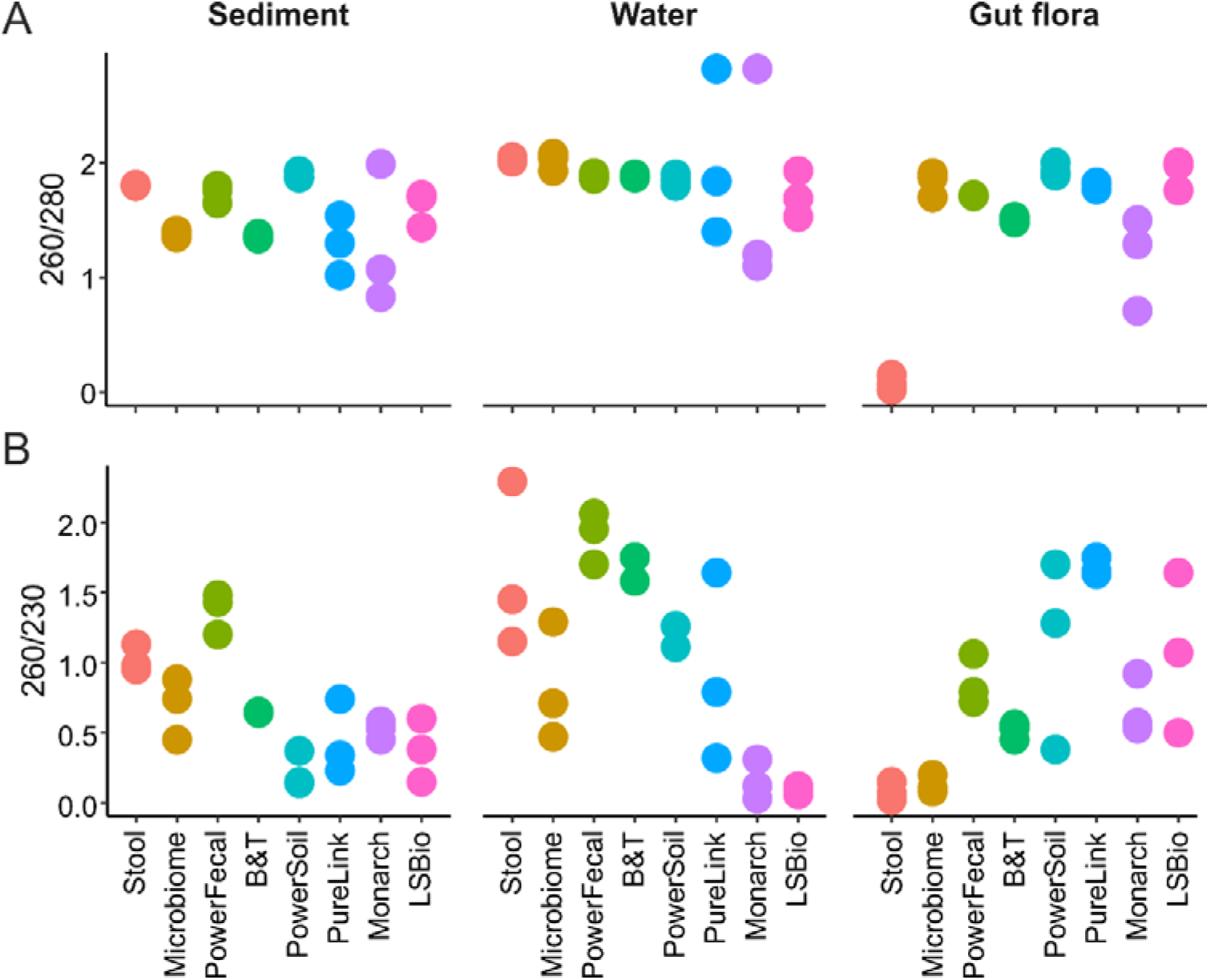
DNA purity assessed by 260/280 (**A**) and 260/230 (**B**) absorption ratios. Data for three technical replicates are shown.

**Fig. S5.**
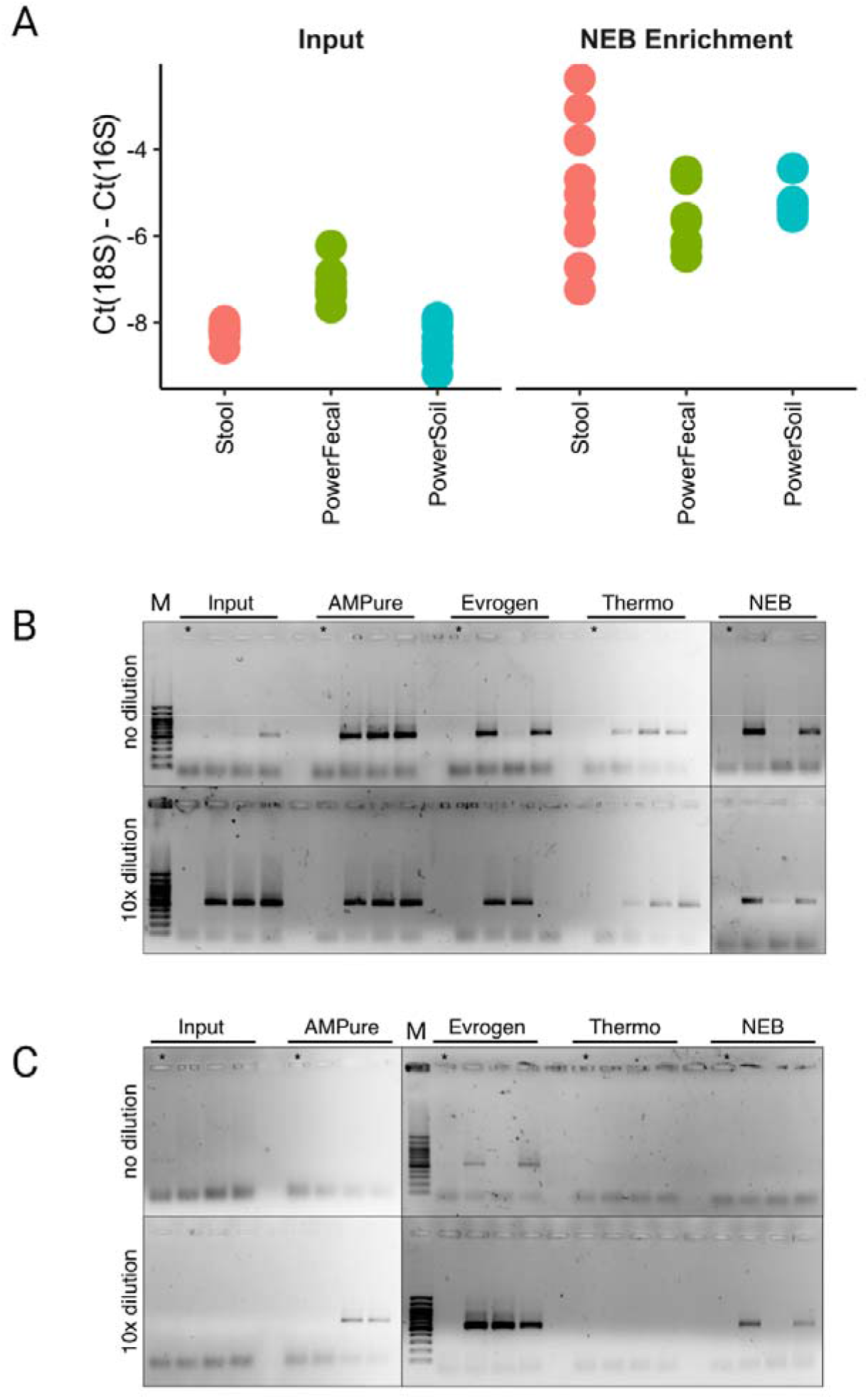
(**A**) Retention of DNA relative to the input amount (50 ng for Stool or 150 ng for Microbiome kits) after additional re-purification procedures. (**B-C**) Results of the 16S rRNA gene PCR with sea sediment samples purified with Stool (**B**) and Microbiome (**C**) kits. PCR was performed with non-diluted and 10-fold diluted input DNA or DNA additionally re-purified with indicated kits (see Methods). *- no-input control that was re-purified in parallel with experimental samples to estimate potential contamination of the re-purification kit solutions with microbial DNA. M – 100 bp Plus GeneRuler DNA ladder (Thermo Scientific). Products were loaded on 3 agarose gels and run in parallel, fragments of different gels are separated by a black line. Uncropped gels can be found in Supplementary Figure 1.

**Fig. S6.**
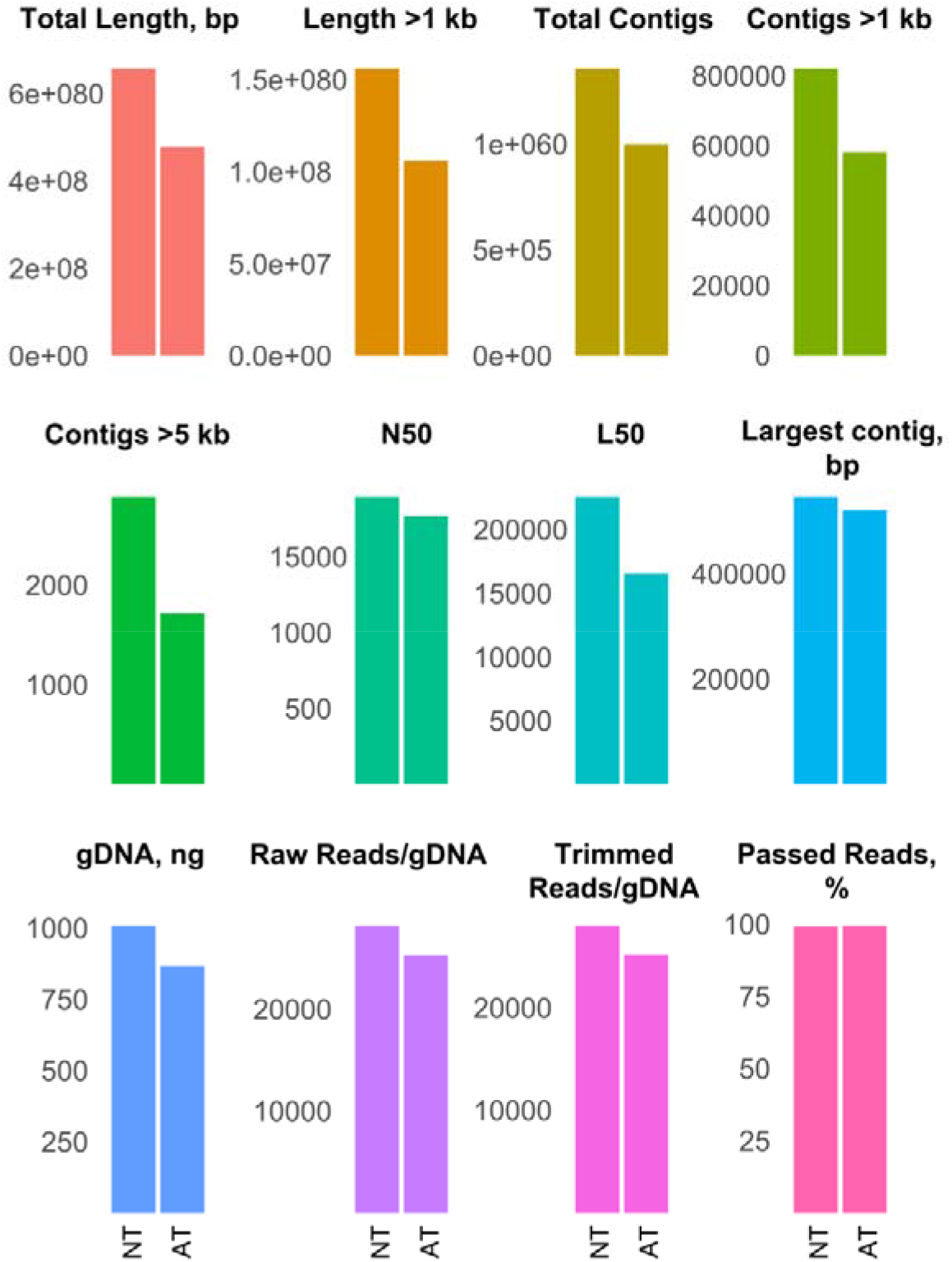
Parameters of shotgun BGI libraries obtained from sediment sample purified with PowerFecal kit (NT – no treatment) and additionally re-purified with Evrogen column kit (AT - additional treatment).

**Fig. S7.**
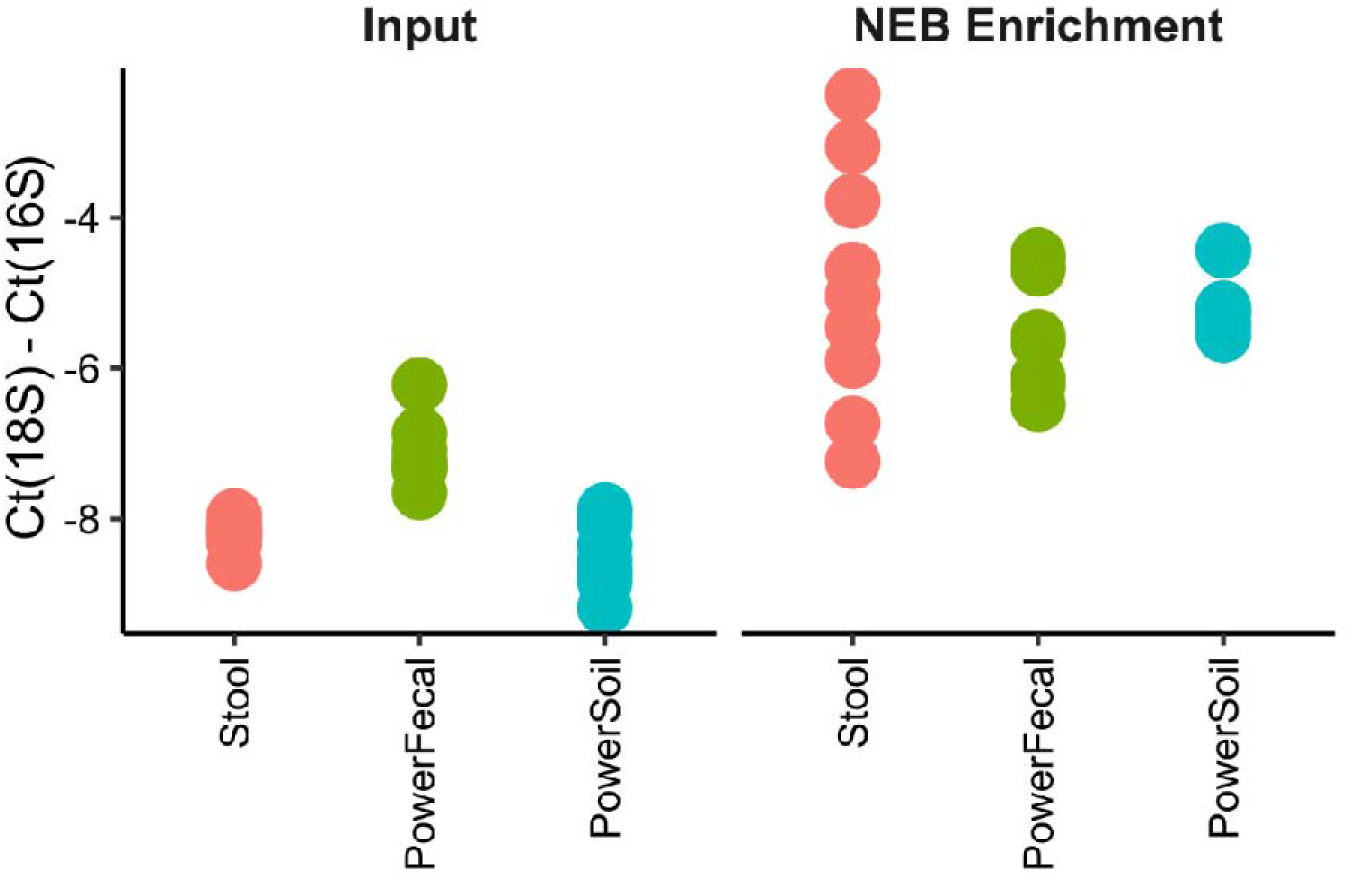
qPCR Ct values obtained with 18S and 16S rRNA gene-specific primers with the input and NEB Enrichment re-purified DNA samples. Data for three technical qPCR replicates for each of the three kit purification replicates are shown.

**Fig. S8.**
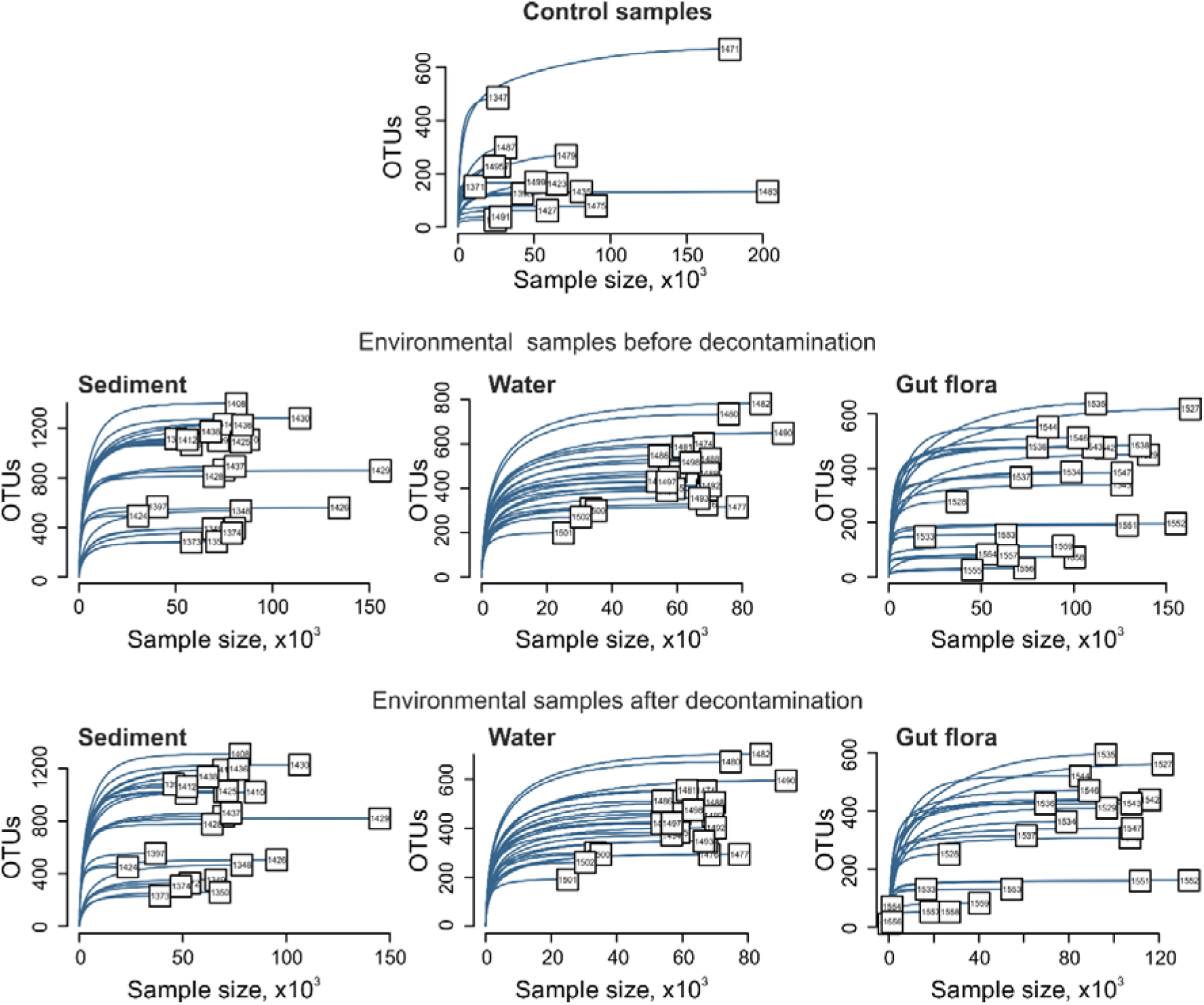
Rarefaction curves for control samples (upper row), natural samples before decontamination (middle), and natural samples after decontamination (bottom row).

**Fig. S9.**
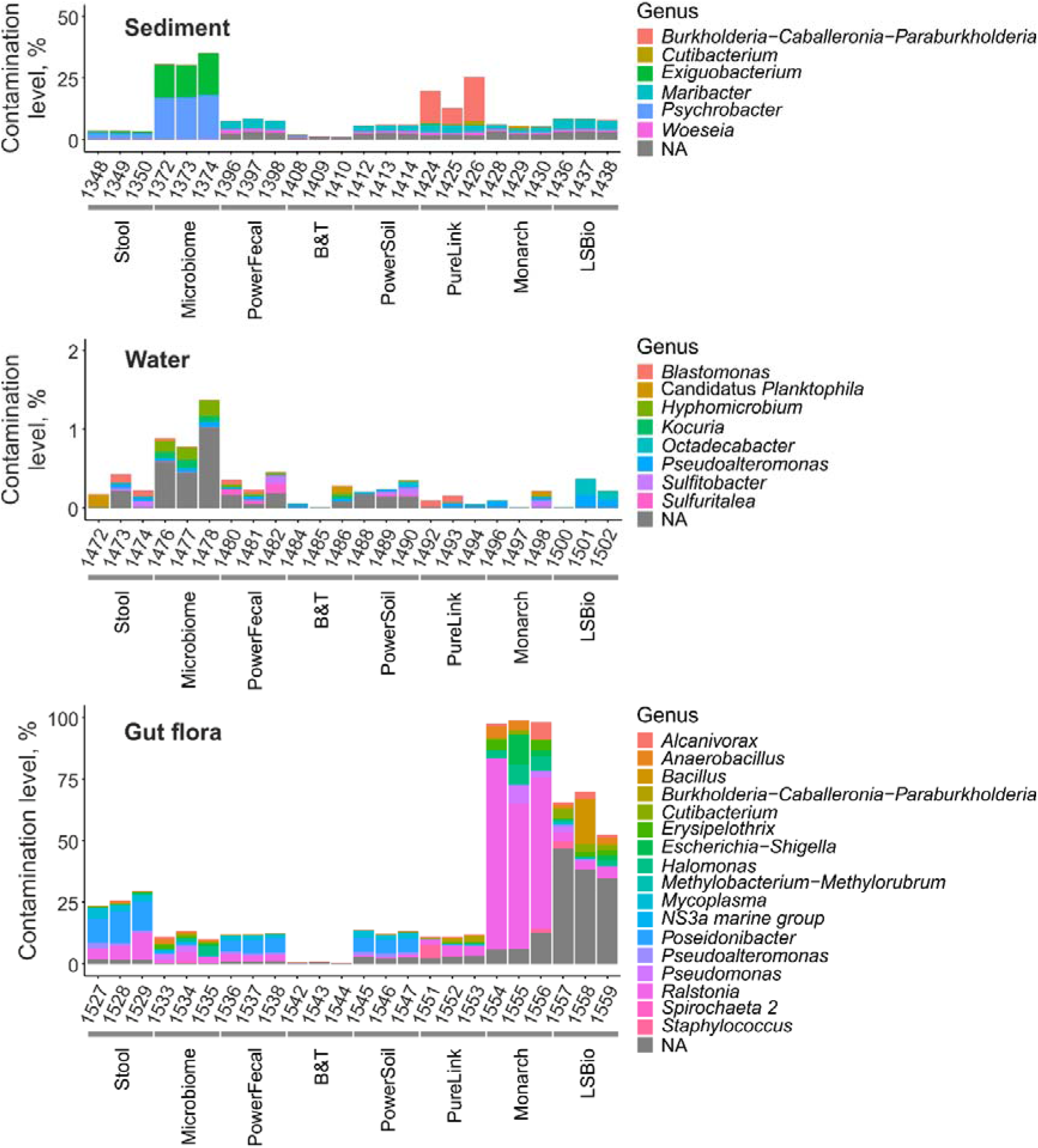
Contamination levels of natural samples. Data is shown for all technical replicates independently. Genera with relative abundances >1% are shown.

**Fig. S10.**
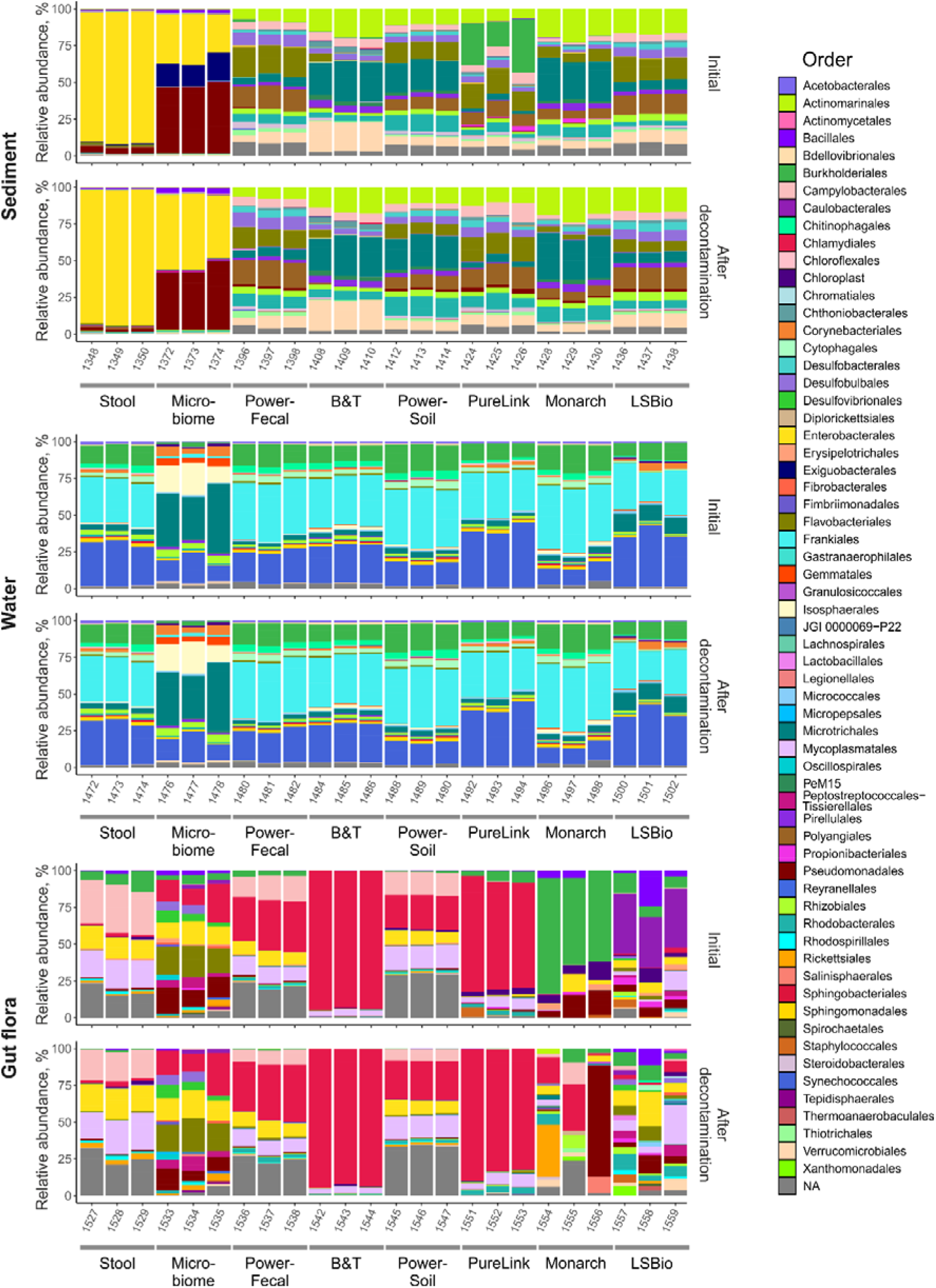
Microbial communities’ composition of natural samples before decontamination (Initial) and after the decontamination on an order level. Data is shown for all technical replicates independently. Orders with relative abundances >1% are shown.

**Fig. S11.**
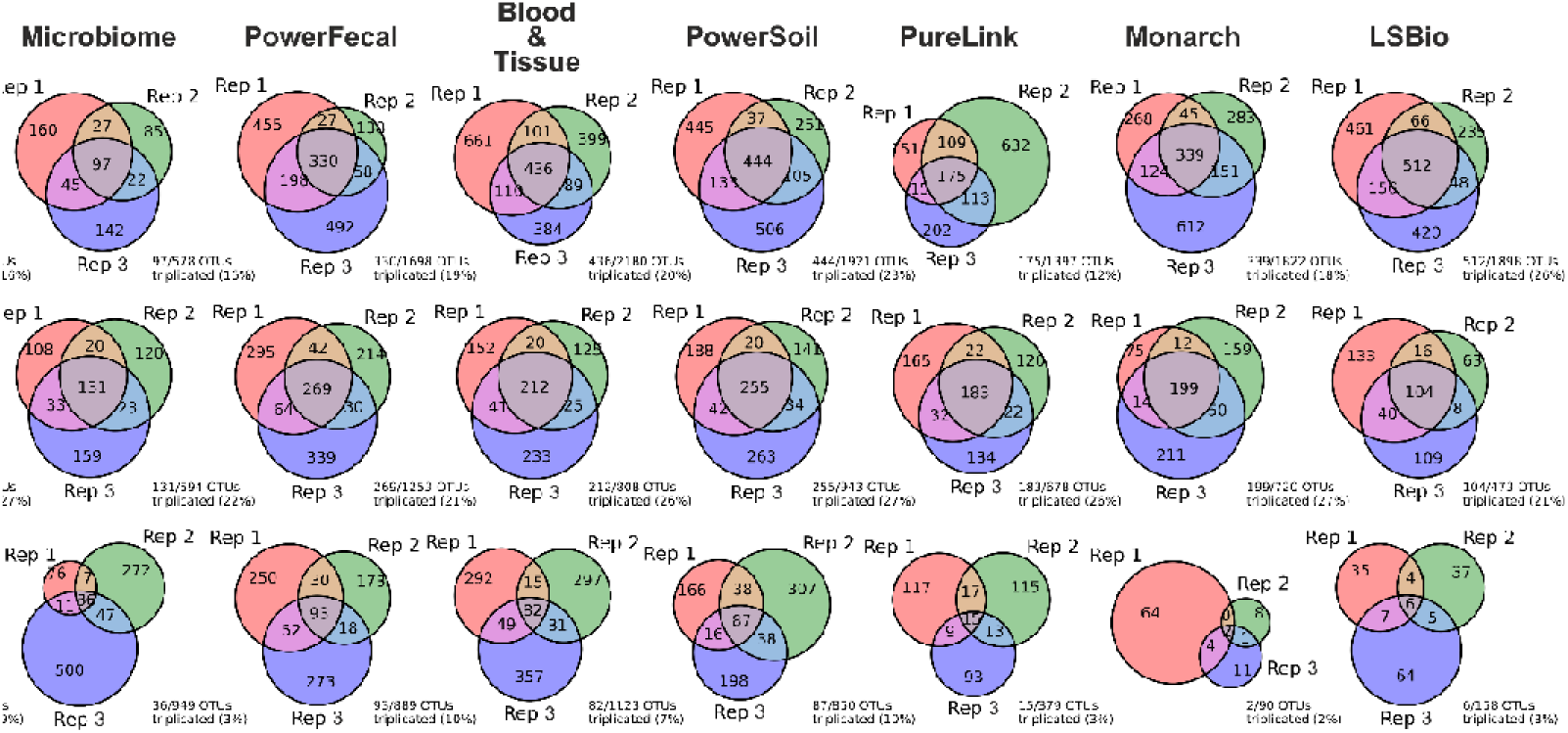
Reproducibility of DNA-extraction kits. Venn diagrams representing the intersections of lists of non-zero OTUs (OTUs with a non-zero abundance) for three technical replicates obtained with specified DNA-extraction kits. Below the diagram, reproducibility level is shown (%) as a fraction of non-zero OTUs found in all three replicates from the total number of unique non-zero OTUs found in at least one replicate.

**Fig. S12.**
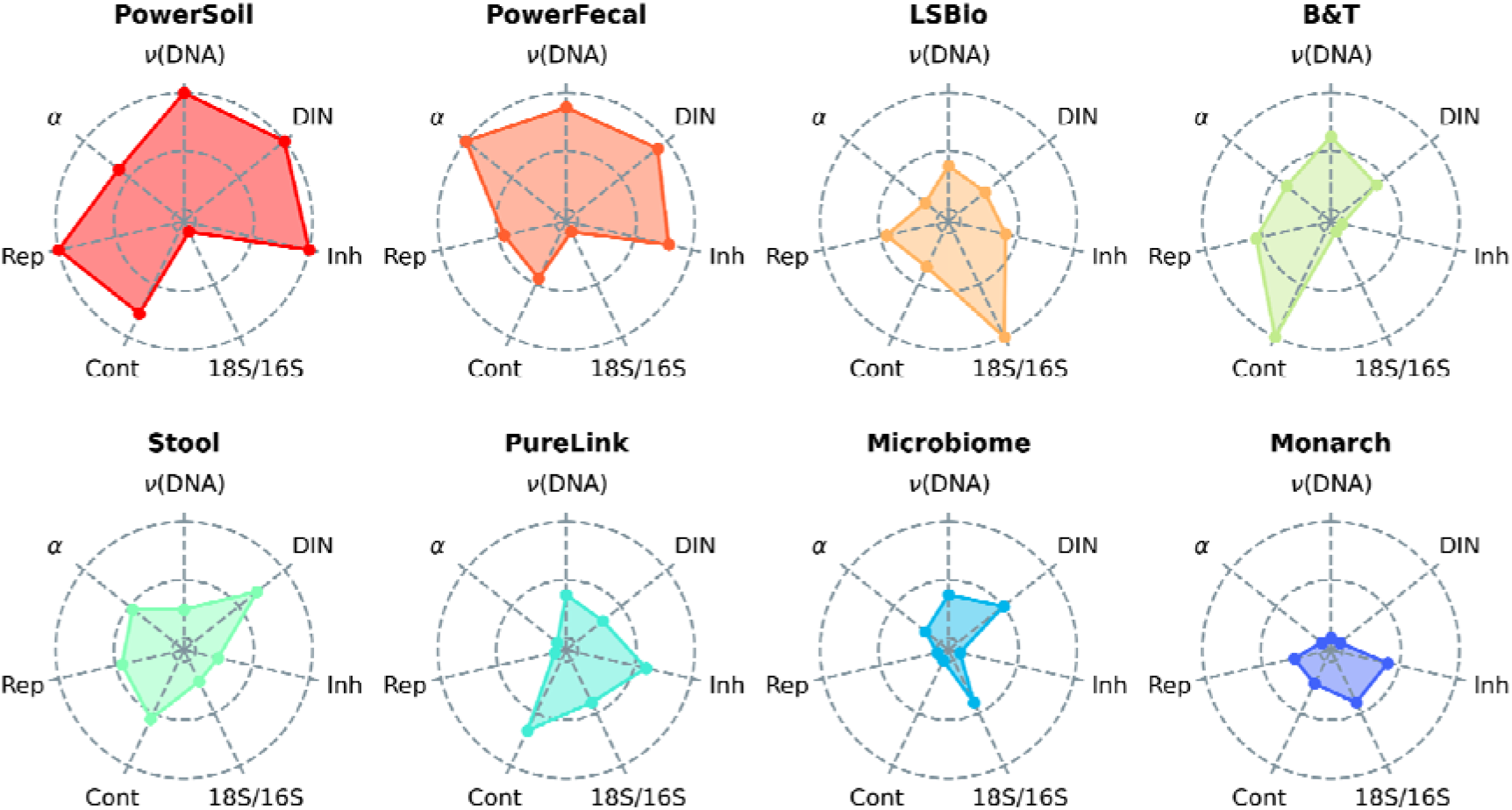
Radar-plots demonstrating the average performance of DNA-extraction kits. LJ(DNA) - DNA yield, DIN - DNA integrity, Inh - presence of PCR inhibitors (higher rank indicates the lower level of inhibitors), 18S/16S - 18S/16S ratio (higher rank indicates the lower ratio), Cont - contamination level (higher rank indicates the lower level of contamination), Rep - reproducibility level, LJ - alpha-diversity. Kits were ordered by the sum of ranks.

**Supplementary Figure 1.**
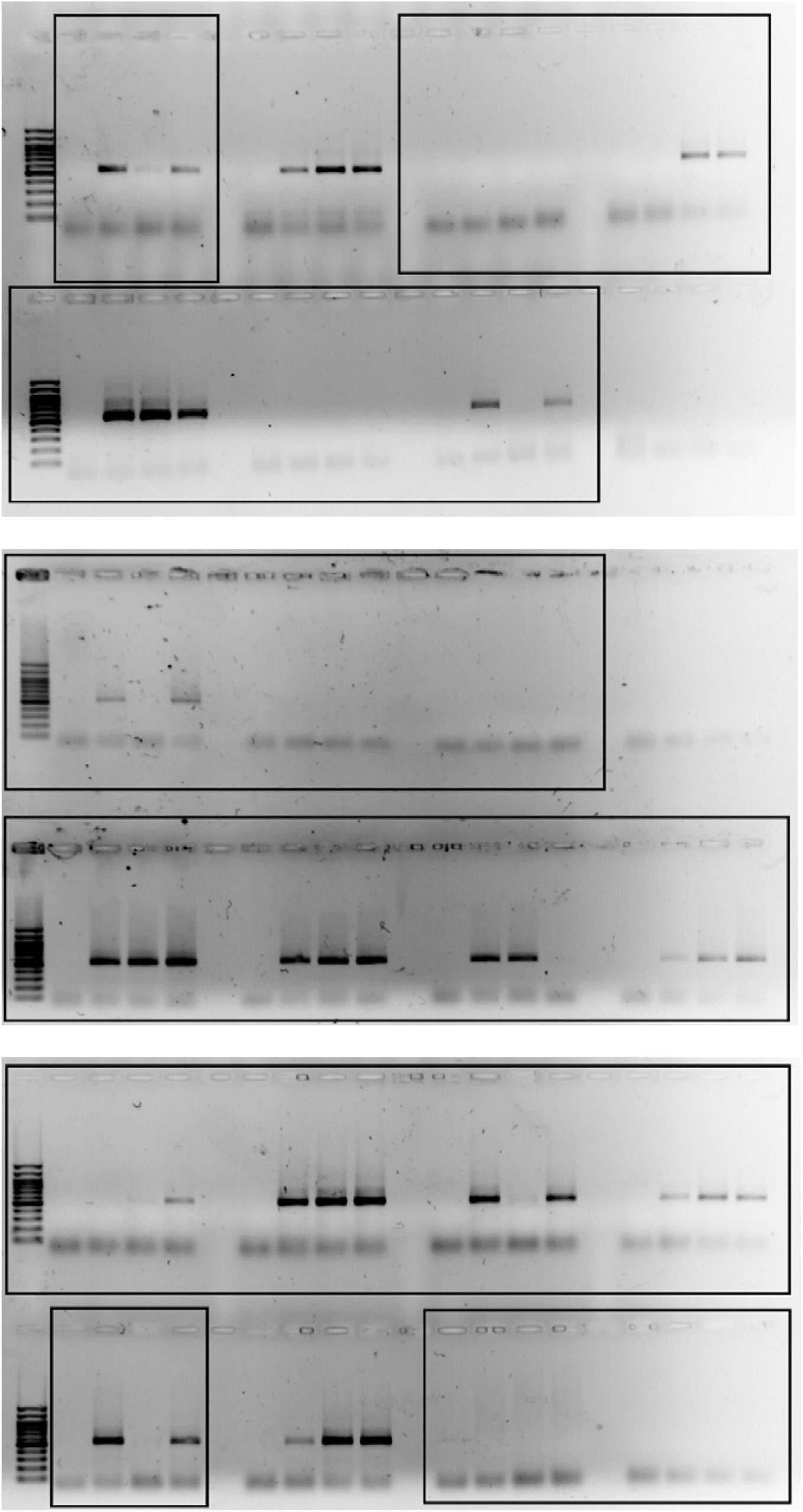
Uncropped agarose gels used to generate Fig. S5B,C. Cropped fragments used to build Fig. S5B,C are shown in frames.

